# Benchmarking Machine Learning Methods for Synthetic Lethality Prediction in Cancer

**DOI:** 10.1101/2023.11.21.567162

**Authors:** Yimiao Feng, Yahui Long, He Wang, Yang Ouyang, Quan Li, Min Wu, Jie Zheng

## Abstract

Synthetic lethality (SL) is a type of genetic interaction that occurs when defects in two genes cause cell death, while a defect in a single gene does not. Targeting an SL partner of a gene mutated in cancer can selectively kill tumor cells. Traditional wet-lab experiments for SL screening are resource-intensive. Hence, many computational methods have been developed for virtual screening of SL gene pairs. This study benchmarks recent machine learning methods for SL prediction, including three matrix factorization and eight deep learning models. We scrutinize model performance using various data splitting scenarios, negative sample ratios, and negative sampling methods on both classification and ranking tasks to assess the models’ generalizability and robustness. Our benchmark analyzed performance differences among the models and emphasized the importance of data and real-world scenarios. Finally, we suggest future directions to improve machine learning methods for SL discovery in terms of predictive power and interpretability.

## Introduction

The synthetic lethal (SL) interaction between genes was first discovered in *Drosophila Melanogaster* about a century ago^1,2^. SL occurs if mutations in two genes result in cell death, but a mutation in either gene alone does not. Based on this observation, Hartwell et al.^3^ and Kealin^4^ suggested that SL could be used to identify new targets for cancer therapy. In the context of cancer, where multiple genes are often mutated, identifying the SL partners of these genes and interfering with their function can lead to cancer cell death, but spare normal cells. PARP inhibitors (PARPi) are the first clinically approved drugs designed by exploiting SL^5^, which target the PARP proteins responsible for DNA repair for the treatment of tumors with BRCA1/2 mutations^5–9^. Clinical trials have shown that PARPi show promising results against tumors such as lung, ovarian, breast, and prostate cancers^9–12^. Despite the success of PARPi, there are still few SL-based drugs that have passed the clinical trials so far, partly due to the lack of techniques to efficiently identify clinically relevant and robust SL gene pairs.

Many methods have been proposed for identifying potential SL gene pairs in the last decade. Various wet-lab experimental methods such as drug screening^13^, RNAi screening^14^, and CRISPR/Cas9 screening^15^ have been used to screen gene pairs with SL relationships^16^. However, due to the large number of pairwise gene combinations (approximately 200 million in human cells)^17^, it is impractical to screen all potential SL pairs by these wet-lab methods. To reduce the search space for SL gene pairs, many computational methods have been proposed. Statistical methods identify SL gene pairs based on hypotheses derived from specific biological knowledge. These methods are generally interpretable because they can reveal statistical patterns between gene pairs^18^. So far, biologists have mainly relied on methods such as DAISY^19^, ISLE^20^, MiSL^21^, SiLi^22^, etc. Additionally, random forests (RF)^23^, a traditional machine learning method, is also frequently used^24–26^, this may also due to its easier to understand nature relative to deep learning. However, the accuracy of these models largely depends on the reliability of the assumptions and the reliability of feature extraction, and they often hard to be scaled up. In contrast, deep learning methods can better capture the complex nonlinear relationships between input and output, enabling them to identify complex patterns in the data. However, deep learning methods have not been well-received by the biological community, largely due to their black-box nature. Interpretability is crucial to whether biologists to adopt deep learning methods.

Two recent reviews on SL^18,27^have comprehensively summarized the data resources as well as computational methods associated with SL; however, neither of them has been able to provide a more comprehensive and systematic assessment of the performance of these methods. In this work, we systematically evaluate the performance of machine learning-based SL prediction methods on different tasks and scenarios, and test the impact of negative samples from different strategies on model performance.

In this study, we perform a comprehensive benchmarking of machine learning methods for SL prediction, including deep learning, and provide some guidelines to biologists regarding the use of the models. First, we collated recently published traditional machine learning and deep learning methods for predicting SL interactions (see Supplementary Table S1 and S2), from which we selected three matrix factorization methods and eight deep learning methods for benchmarking (Table 3). To standardize input and output formats and to facilitate data processing and result aggregation, we preprocessed SL labels and related data from sources such as SynLethDB^28^ (SynLethKG), Gene Ontology^29^ (GO), BioGRID^30^ (PPI), and KEGG^31^ (pathways). To assess the generalizability and robustness of the models, we incorporated three data segmentation methods (DSMs) and four positive-to-negative ratios (PNRs) into our experimental design. Additionally, we investigated the impact of negative sample quality on the model performance by utilizing three negative sampling methods (NSMs). Finally, we also performed two prediction tasks (i.e., classification and ranking) to identify the most probable SL gene pairs. Benchmarking results indicate that integrating information from multiple data sources is beneficial for predicting results, but as an ideal data source, the utilization of KG still needs to be explored. Additionally, our study extends the SL prediction problem to a more realistic scenario, provides valuable insights into the performance of different AI approaches to SL prediction, and further provides some suggestions for future development of SL prediction methods.

## Results

### Benchmarking pipeline

To evaluate machine learning methods for predicting SL interactions, we selected 11 methods published in recent years, including three matrix factorization-based methods (SL^2^MF^32^, CMFW^33^, and GRSMF^34^) and eight graph neural network-based methods (DDGCN^35^, GCATSL^36^, SLMGAE^37^, MGE4SL^38^, PTGNN^39^, KG4SL^40^, PiLSL^41^, and NSF4SL^42^), see Table 3 for more details. The input data for these models varies, and in addition to SL labels, many kinds of data are used to predict SL, including GO^29^, PPI^30^, pathways^31^, and KG^28^, etc., and the detailed data requirements for each model are shown in Table 4. We believe that how many kinds of data inputs a model can accept is a function of the model’s own capabilities, and we focus on the model’s performance in various scenarios. To accomplish this, we designed 36 experimental scenarios, taking into account 3 different data splitting methods (DSMs), 4 positive and negative sample ratios (PNRs), and 3 negative sampling methods (NSMs) as shown in Figure 1. In particular, these scenarios can be described as: (NSM_N_, CV_*i*_, 1:R), where N∈ {Rand, Exp, Dep}; *i* ∈{1, 2, 3} ; R*∈* {1, 5, 20, 50} (see Methods for specific settings). After obtaining the results of all the methods in various experimental scenarios, we evaluated their performance for both classification and ranking tasks, and designed an overall score (see Methods) to better quantify their performance in Figure 2. We also evaluated the scalability of all the models, including their computational efficiency and code quality. For a more visual presentation of the benchmarking results, we provided figures and tables (Figure 2, Table 1, 2 and Supplementary Materials Figs. S1-S9) to show the results under various scenarios.

**Table 1.**
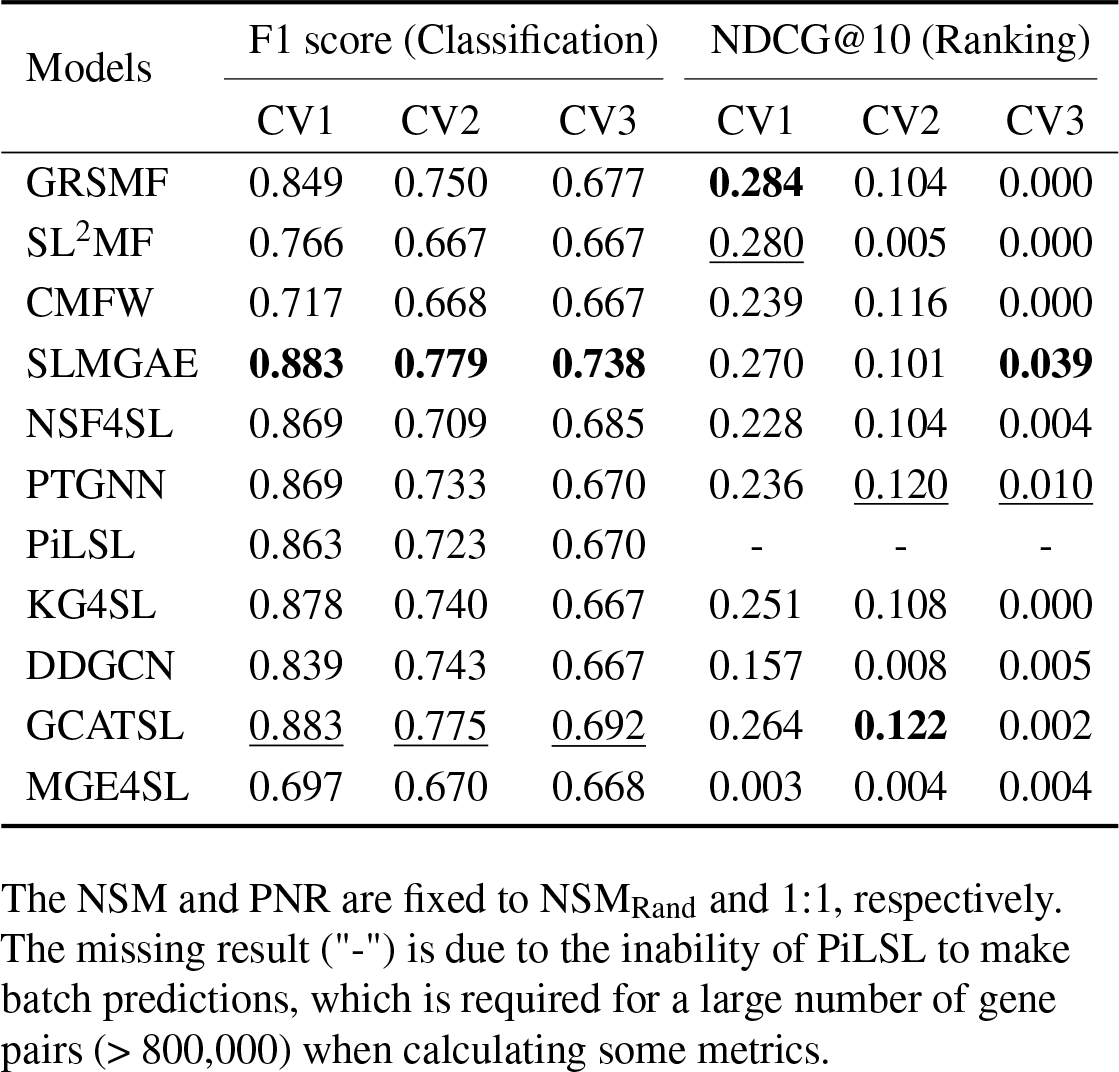
The performance of the models under different DSMs.

| Models | F1 score (Classification) |  |  | NDCG@10 (Ranking) |  |  |
| --- | --- | --- | --- | --- | --- | --- |
|  | CV1 | CV2 | CV3 | CV1 | CV2 | CV3 |
| GRSMF | 0.849 | 0.750 | 0.677 | **0.284** | 0.104 | 0.000 |
| SL<sup>2</sup>MF | 0.766 | 0.667 | 0.667 | <u>0.280</u> | 0.005 | 0.000 |
| CMFW | 0.717 | 0.668 | 0.667 | 0.239 | 0.116 | 0.000 |
| SLMGAE | **0.883** | **0.779** | **0.738** | 0.270 | 0.101 | **0.039** |
| NSF4SL | 0.869 | 0.709 | 0.685 | 0.228 | 0.104 | 0.004 |
| PTGNN | 0.869 | 0.733 | 0.670 | 0.236 | <u>0.120</u> | <u>0.010</u> |
| PiLSL | 0.863 | 0.723 | 0.670 | - | - | - |
| KG4SL | 0.878 | 0.740 | 0.667 | 0.251 | 0.108 | 0.000 |
| DDGCN | 0.839 | 0.743 | 0.667 | 0.157 | 0.008 | 0.005 |
| GCATSL | <u>0.883</u> | <u>0.775</u> | <u>0.692</u> | 0.264 | **0.122** | 0.002 |
| MGE4SL | 0.697 | 0.670 | 0.668 | 0.003 | 0.004 | 0.004 |

The NSM and PNR are fixed to NSM<sub>Rand</sub> and 1:1, respectively. The missing result ("-") is due to the inability of PiLSL to make batch predictions, which is required for a large number of gene pairs (> 800,000) when calculating some metrics.

**Table 2.**
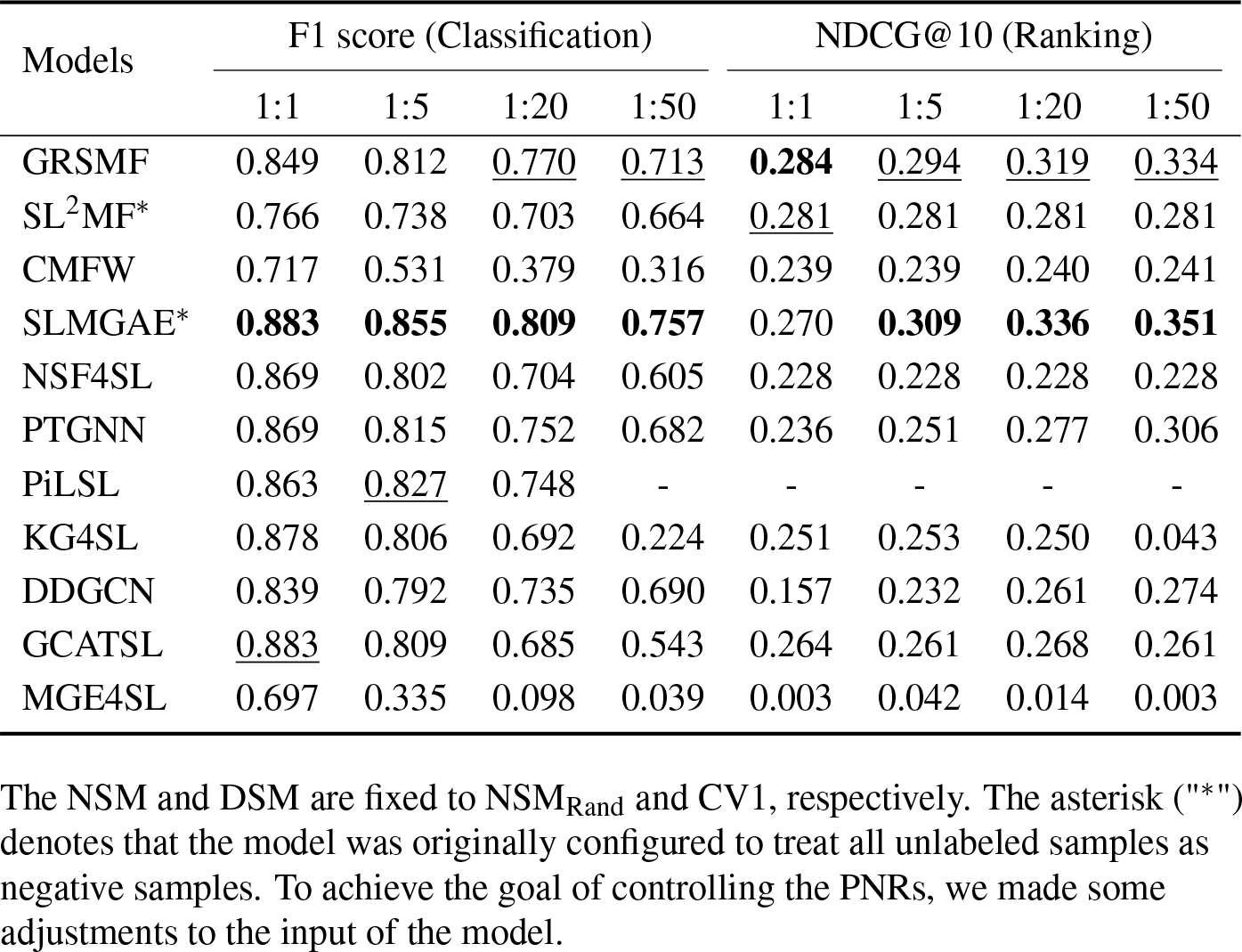
The performance of the models under different PNRs.

| Models | F1 score (Classification) |  |  |  | NDCG@10 (Ranking) |  |  |  |
| --- | --- | --- | --- | --- | --- | --- | --- | --- |
|  | 1:1 | 1:5 | 1:20 | 1:50 | 1:1 | 1:5 | 1:20 | 1:50 |
| GRSMF | 0.849 | 0.812 | <u>0.770</u> | <u>0.713</u> | **0.284** | <u>0.294</u> | <u>0.319</u> | <u>0.334</u> |
| SL<sup>2</sup>MF* | 0.766 | 0.738 | 0.703 | 0.664 | <u>0.281</u> | 0.281 | 0.281 | 0.281 |
| CMFW | 0.717 | 0.531 | 0.379 | 0.316 | 0.239 | 0.239 | 0.240 | 0.241 |
| SLMGAE* | **0.883** | **0.855** | **0.809** | **0.757** | 0.270 | **0.309** | **0.336** | **0.351** |
| NSF4SL | 0.869 | 0.802 | 0.704 | 0.605 | 0.228 | 0.228 | 0.228 | 0.228 |
| PTGNN | 0.869 | 0.815 | 0.752 | 0.682 | 0.236 | 0.251 | 0.277 | 0.306 |
| PiLSL | 0.863 | <u>0.827</u> | 0.748 | - | - | - | - | - |
| KG4SL | 0.878 | 0.806 | 0.692 | 0.224 | 0.251 | 0.253 | 0.250 | 0.043 |
| DDGCN | 0.839 | 0.792 | 0.735 | 0.690 | 0.157 | 0.232 | 0.261 | 0.274 |
| GCATSL | <u>0.883</u> | 0.809 | 0.685 | 0.543 | 0.264 | 0.261 | 0.268 | 0.261 |
| MGE4SL | 0.697 | 0.335 | 0.098 | 0.039 | 0.003 | 0.042 | 0.014 | 0.003 |

The NSM and DSM are fixed to NSM<sub>Rand</sub> and CV1, respectively. The asterisk ("\*") denotes that the model was originally configured to treat all unlabeled samples as negative samples. To achieve the goal of controlling the PNRs, we made some adjustments to the input of the model.

**Table 3.**
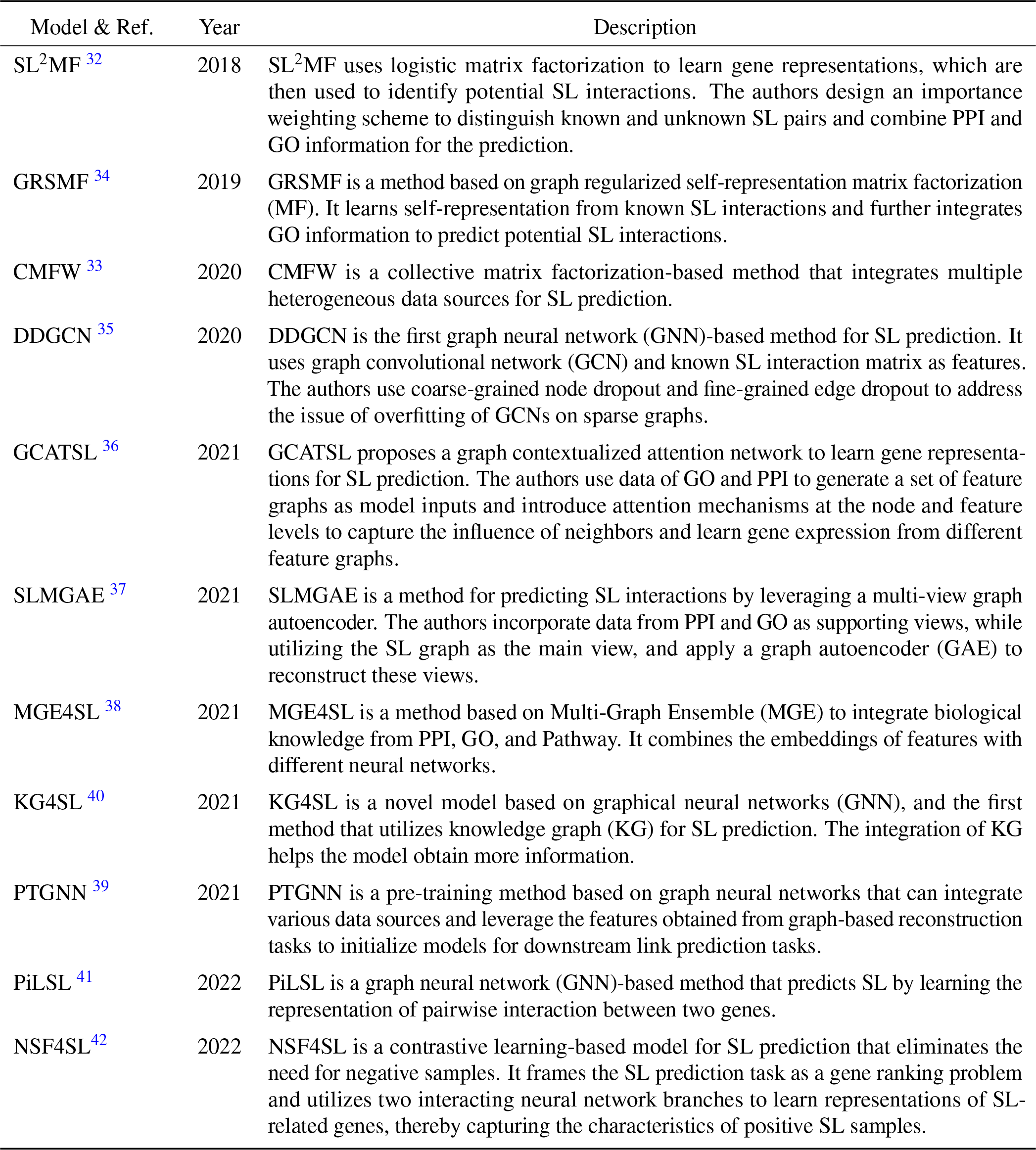
List of supervised machine learning method for SL prediction.

**Table 4.**
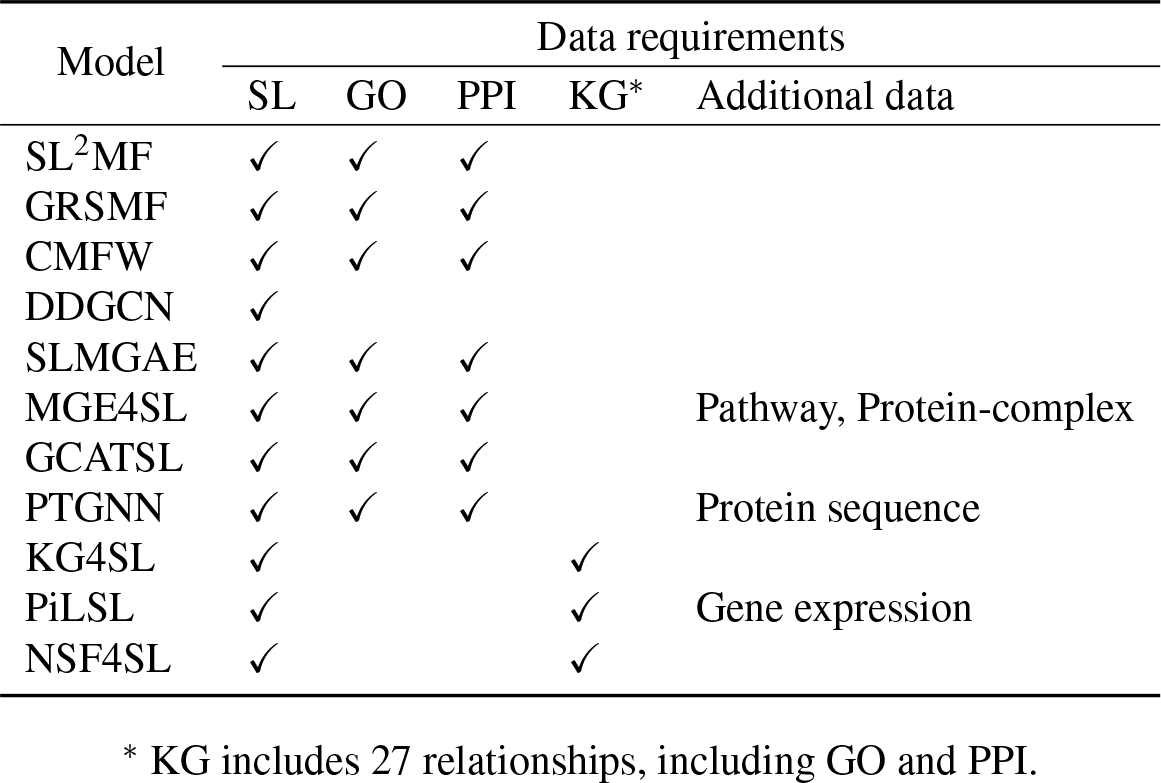
Data requirements of all models compared in this work.

| Model | Data requirements |  |  |  |  |
| --- | --- | --- | --- | --- | --- |
|  | SL | GO | PPI | KG* | Additional data |
| SL<sup>2</sup>MF | ✓ | ✓ | ✓ |  |  |
| GRSMF | ✓ | ✓ | ✓ |  |  |
| CMFW | ✓ | ✓ | ✓ |  |  |
| DDGCN | ✓ |  |  |  |  |
| SLMGAE | ✓ | ✓ | ✓ |  |  |
| MGE4SL | ✓ | ✓ | ✓ |  | Pathway, Protein-complex |
| GCATSL | ✓ | ✓ | ✓ |  |  |
| PTGNN | ✓ | ✓ | ✓ |  | Protein sequence |
| KG4SL | ✓ |  |  | ✓ |  |
| PiLSL | ✓ |  |  | ✓ | Gene expression |
| NSF4SL | ✓ |  |  | ✓ |  |

\* KG includes 27 relationships, including GO and PPI.

**Figure 1.**
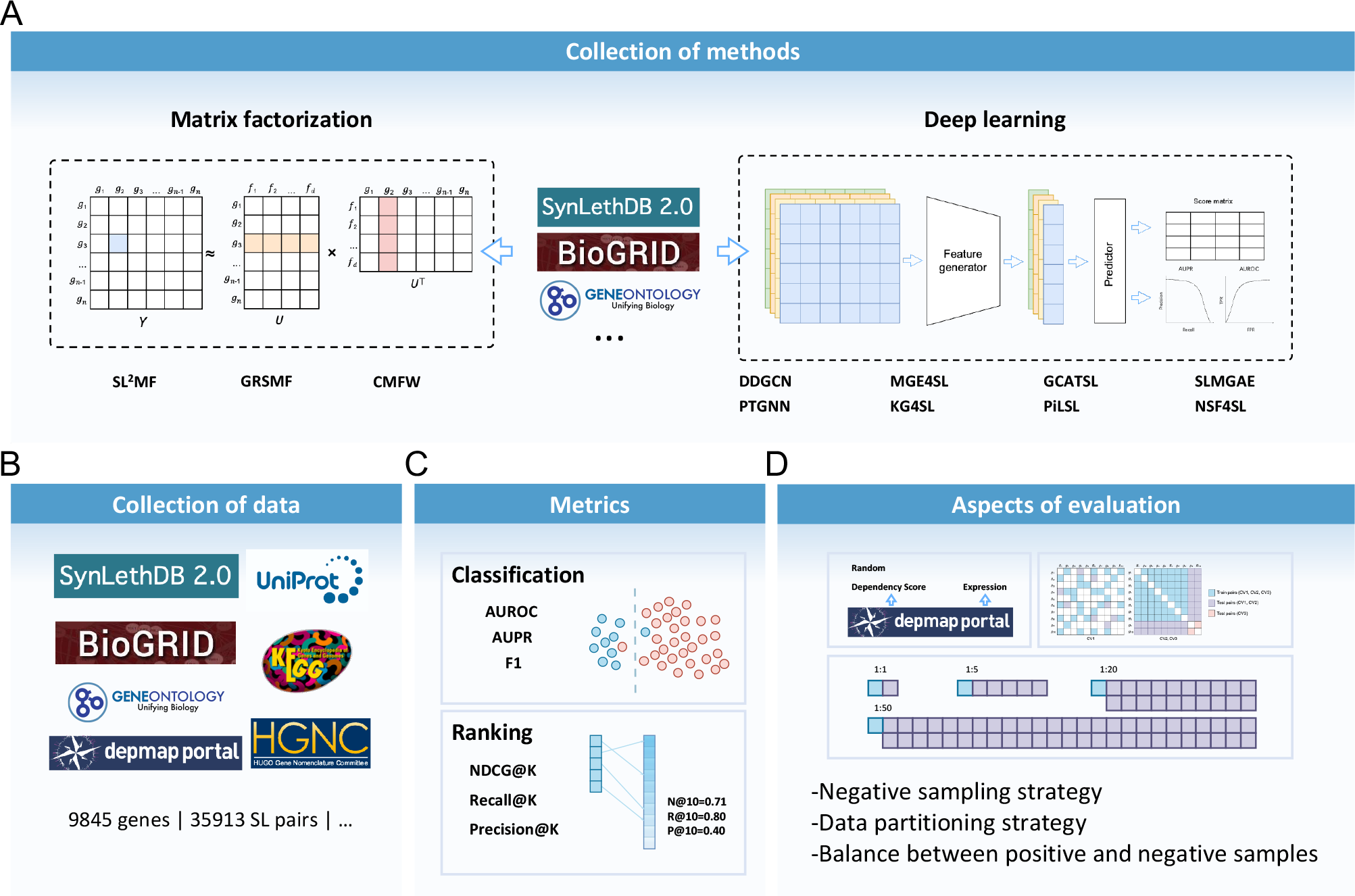
Workflow of the benchmarking study. **A**. A total of eleven methods are compared, including three matrix factorization-based methods and eight deep learning methods. **B**. We collected data from different databases to build a benchmark dataset. **C**. This study compared the performance of the models in both classification and ranking tasks. **D**. We also designed various experimental scenarios, including different negative sampling methods, positive-negative sample ratios, and data partitioning methods. The combinations of these scenarios constitute a task space ranging from simple to difficult tasks.

**Figure 2.**
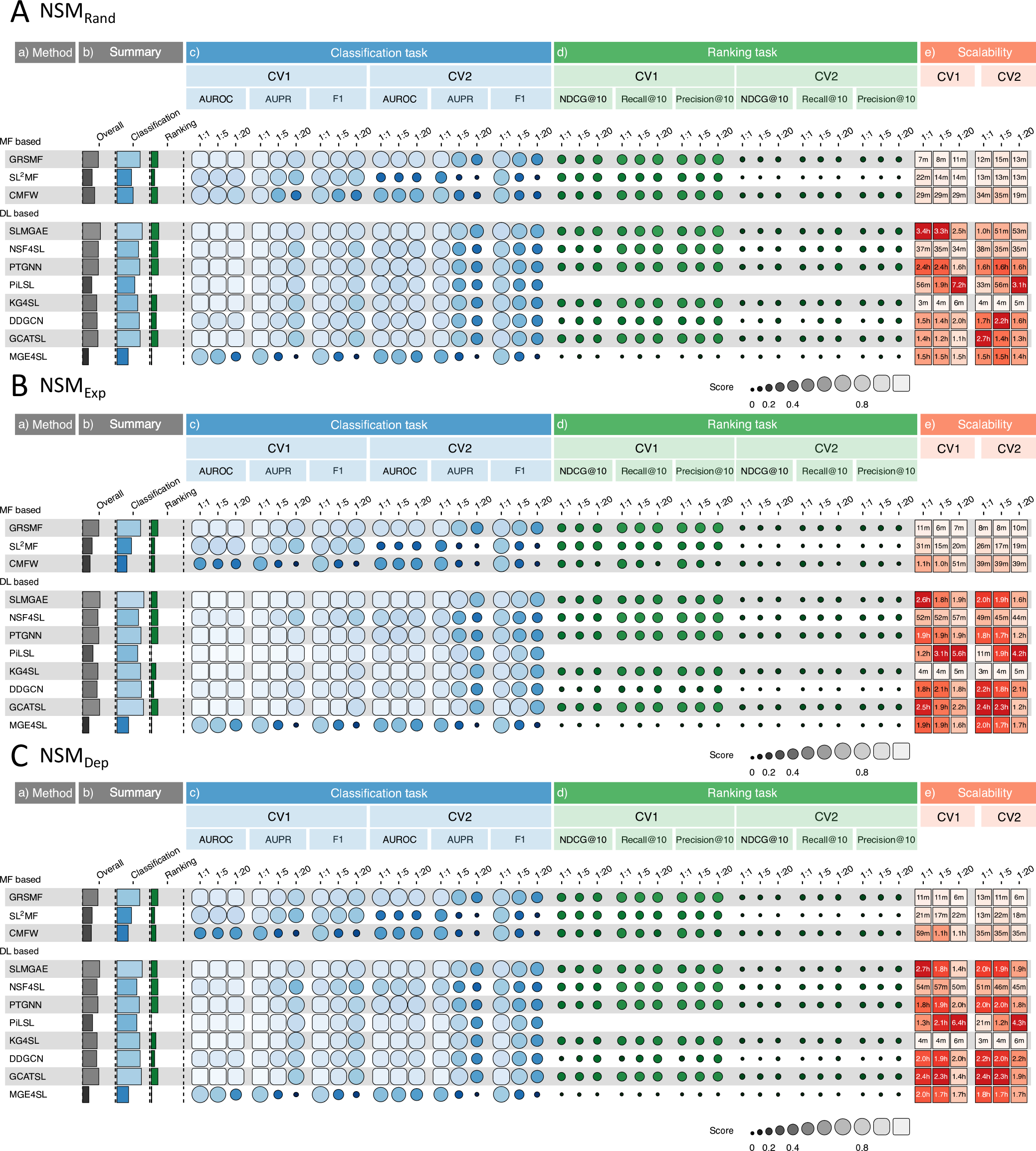
Performance of the models. **A, B** and **C** are the performance of the model under NSM_Rand_, NSM_Exp_ and NSM_Dep_, respectively, where lighter colors indicate better performance. The figure contains five parts of information: a) A list of the 11 models. b) The overall scores of the models and the combined scores under the classification task and the ranking task only. c) and d) The performance of the models under the classification and ranking tasks, including six experimental scenarios consisting of 2 DSMs and 3 PNRs. e) The average time required for the models to complete one cross validation

### Classification and ranking

For the problem of predicting SL interactions, most current methods still consider it as a classification problem, i.e., to determine whether a given gene pair has SL interaction. However, models with classification capabilities alone are insufficient for biologists, who need a curated list of genes that may have SL relationships with the genes they are familiar with. This list can empower biologists to conduct wet-lab experiments such as CRISPR-based screening. Among the evaluated methods, only NSF4SL originally regards this problem as a gene recommendation task, while other methods belong to the traditional discriminative models. To compute metrics for both tasks using these models, we adjusted the output layer, this modification ensures that every model produces a floating point score as its output.

To assess the overall performance of the models in classification and ranking tasks, we employed separate Classification scores and Ranking scores (see Methods). Figure 2 presents these scores and the model’s performance across different scenarios and metrics. Based on the Classification scores, we found that when using negative samples filtered based on NSM_Exp_, the models usually had the best performance for the classification task. Among them, SLMGAE, KG4SL, and PiLSL performed the best with Classification scores of 0.854, 0.853, and 0.851, respectively. On the ranking task, the models performed slightly better under the scenario of NSM_Rand_, and the top three methods were SLMGAE, PTGNN, and GRSMF, achieved Ranking scores of 0.233, 0.222, and 0.212, respectively. From these scores, SLMGAE is the model with the best overall performance.

In the following three sections, for the purpose of consistently assessing the performance of each model across classification and ranking tasks, we have designated a single metric for each task. Given our focus on the accurate classification of positive samples and the imbalance between positive and negative samples in experimental settings, we primarily employ F1 scores to gauge the models’ classification performance. Additionally, to appraise the model’s effectiveness in the ranking task, we mainly rely on the NDCG@10 metric, which takes into account the relevance and ranking of the genes in the SL prediction list.

### Generalizability to unseen genes

In this section, we utilized three different data splitting methods (DSMs) for cross validation, namely CV1, CV2, and CV3 (see Methods), in the order of increasing difficulty. The performance of models given these DSMs reflects their ability to generalize from known to unknown SL relationships.

Among the three DSMs, CV1 is the most frequently used method for cross validation. However, this method exclusively provides accurate predictions for genes present in the training set, lacking the ability to extend its predictive capabilities to genes unseen during training. The CV2 scenario can be characterized as a semi-cold start problem, i.e., one and only one gene in a gene pair is present in the training set. This scenario holds significant practical implications. Considering the existence of ∼10,000 known genes involved in SL interactions, there is a substantial number of human genes remain unexplored. These genes, which have not yet received enough attention, likely include numerous novel SL partner genes of known primary genes mutated in cancers. CV3 is a complete cold-start problem, i.e., nether of the two genes is in the training set. Under CV3, the model must adeptly discern common patterns of SL relationships, to generalize to genes not encountered during training.

For the convenience of discussion, we fixed NSM to NSM_Rand_ and PNR to 1:1, i.e., our scenario is (NSM_Rand_, CV_*i*_, 1:1), where *i* = 1, 2, 3. See Supplementary Data 1, 2, and 3 for the complete results under all scenarios.

Table 1 shows the performance of the models under different DSMs while NSM and PNR are fixed to NSM_Rand_ and 1:1, respectively. From the table, it can be seen that, for the classification task, SLMGAE, GCATSL, and KG4SL performed better.

under the CV1 scenario, with F1 scores greater than 0.877. By contrast, CMFW and MGE4SL performed poorly, with F1 score less than 0.720. Under CV2, all the selected methods showed significant performance degradation compared to CV1. For example, the F1 scores of SLMGAE and GCATSL, still the top two methods, dropped to 0.779 and 0.775, respectively, while both of CMFW and MGE4SL decreased to less than 0.670. When the DSM is changed to CV3, only the F1 score of SLMGAE can still be above 0.730, while all the other methods drop below 0.700. For the ranking task, GRSMF, SL^2^MF and SLMGAE exhibited better performance under the CV1 scenario with NDCG@10 greater than 0.270. However, under CV2, the NDCG@10 of almost all methods except for GCATSL and PTGNN are lower than 0.120. Lastly, when the DSM is CV3, the NDCG@10 scores of all the methods in this scenario becomes very low (lower than 0.010) except SLMGAE. Generally, SLMGAE, GCATSL, and GRSMF have good generalization capabilities. In addition, CV3 is a highly challenging scenario for all models, especially for the ranking task.

Moreover, Figure 3 displays the predicted score distributions of gene pairs in the training and testing sets for the SLMGAE and GCATSL models, respectively (see all methods in Supplementary Materials Figs. S10-S20). It is noteworthy that when the DSM is CV1, both models can differentiate between positive and negative samples effectively. As the challenge of generalization increases (in the case of CV2), there is a considerable change in the distribution of sample scores in the two models. In particular, for GCATSL, almost all negative sample scores in this model are concentrated around 0.5, while positive sample scores start to move towards the middle; for SLMGAE, only the positive sample distribution in the test set was significantly affected. When the task scenario becomes the most difficult CV3, the score distribution shift of the positive samples in the test set of the two models is more pronounced. For GCATSL, almost all sample scores in the test set are concentrated around 0.4, and the model cannot fail to have predictive ability in this scenario. Comparatively, SLMGAE is capable of distinguishing the samples in the test set with a relatively high degree of accuracy.

**Figure 3.**
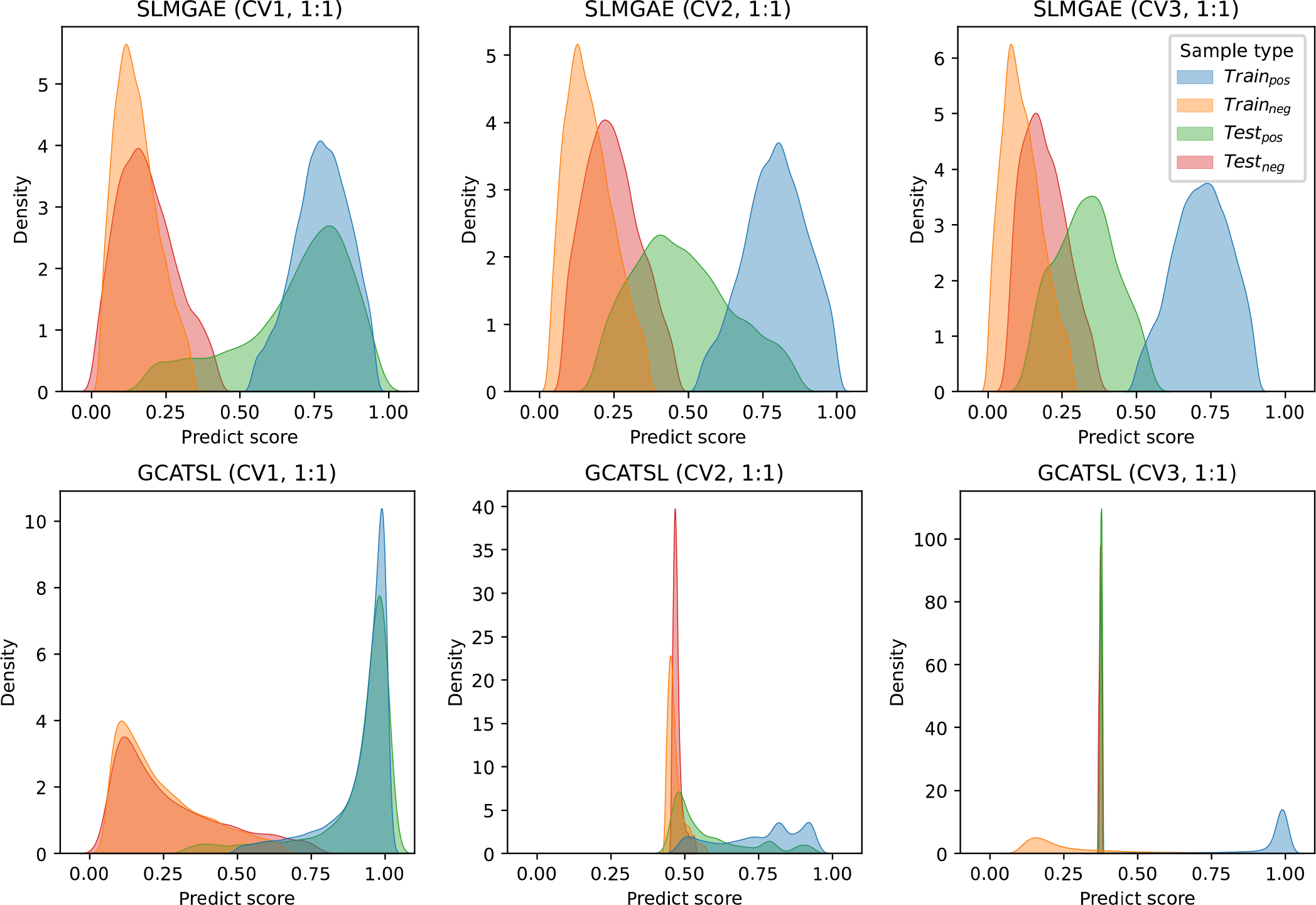
Distribution of predicted score. The scores of gene pairs in the training and testing sets for the SLMGAE and GCATSL models are displayed in the first and second lines, respectively, under various data splitting methods (DSMs). The results are obtained with NSM_Rand_.

### Robustness to increasing number of negative samples

In our study, so far, all negative samples used in training have been screened from unknown samples. As such, there could be false negative samples among the gene pairs categorized as negative. This situation could inadvertently introduce noise into the model’s training process. Furthermore, given the substantial disparity between the numbers of non-SL pairs and SL pairs, these models encounter the issue of imbalanced data. To assess the robustness of these models to noise stemming from negative samples, we conducted experiments by gradually increasing the number of negative samples. In our study, the number of negative samples is set to four levels: equal to the number of positive samples (1:1), five times the number of positive samples (1:5), twenty times the number of positive samples (1:20), and fifty times the number of positive samples (1:50). Notably, the 1:1 ratio corresponds to the conventional experimental configuration frequently adopted.

In this section, our experimental scenario for comparison is denoted as (NSM_Rand_, CV1, 1:R) where R = 1, 5, 20, 50. From Table 2, it can be seen that as the number of negative samples increases, the models’ performance (F1 score) in classification tasks gradually decreases. This phenomenon is particularly pronounced for CMFW and MGE4SL. When the number of negative samples increases from one to five times that of positive samples (PNR is 1:5), the F1 scores of CMFW and MGE4SL drop dramatically from around 0.700 to 0.531 and 0.335, respectively. By contrast, several other methods, namely SLMGAE, PTGNN, PiLSL, KG4SL, and GCATSL, maintain their F1 scores above 0.800. When the number of negative samples is further increased to twenty times the number of positive samples (PNR is 1:20), only SLMGAE achieved an F1 score above 0.800, while CMFW and MGE4SL dropped to 0.379 and 0.098, respectively. Finally, when the number of negative samples is fifty times that of positive samples (PNR is 1:50), SLMGAE still outperformed other models with an F1 score of 0.757, followed by GRSMF with an F1 score of 0.713. Notably, compared with previous PNRs, KG4SL and GCATSL experienced a significant decline in their F1 scores, dropping to 0.224 and 0.543, respectively. On the other hand, in the context of ranking task, when PNR = 1:5, SLMGAE, GRSMF, PTGNN, and DDGCN exhibited a slight improvement in NDCG@10 than PNR is 1:1. At PNR=1:20, the NDCG@10 for SLMGAE, GRSMF, PTGNN, and DDGCN continued to rise. Lastly, the NDCG@10 values for SLMGAE and GRSMF continue to increase to 0.351 and 0.334, respectively, when the PNR is changed to 1:50. Generally, SLMGAE and GRSMF have stronger robustness.

Figure 4 displays the distribution of the scores for positive and negative samples predicted by SLMGAE and GCATSL across different PNRs. The figure show the impact of the number of negative samples on the score of the given gene pair evaluated by the models. As the number of negative samples increases, an increasing number of positive samples in the testing set are assigned lower scores. Despite this effect, the majority of samples are still correctly classified by SLMGAE. But for GCATSL, when the number of negative samples is twenty times that of positive samples, i.e., PNR=1:20, almost all samples are assigned very low scores. When the PNR becomes 1:50, almost all the scores given by GCATSL are concentrated in a very small score range (around 0), and the predictions of the model are no longer reliable. Furthermore, under the results of SLMGAE, it is evident that the distribution of negative samples becomes increasingly concentrated with a higher number of negative samples. And a notable phenomenon is observed in the previous results (Table 2), that certain models exhibit improved performance in the ranking task as the number of negative samples increases. We hypothesize that this improvement is attributed to the higher concentration of scores among negative samples, resulting in a greater number of positive samples achieving higher rankings. Consequently, the performance of some models under ranking tasks improves with increasing PNR.

**Figure 4.**
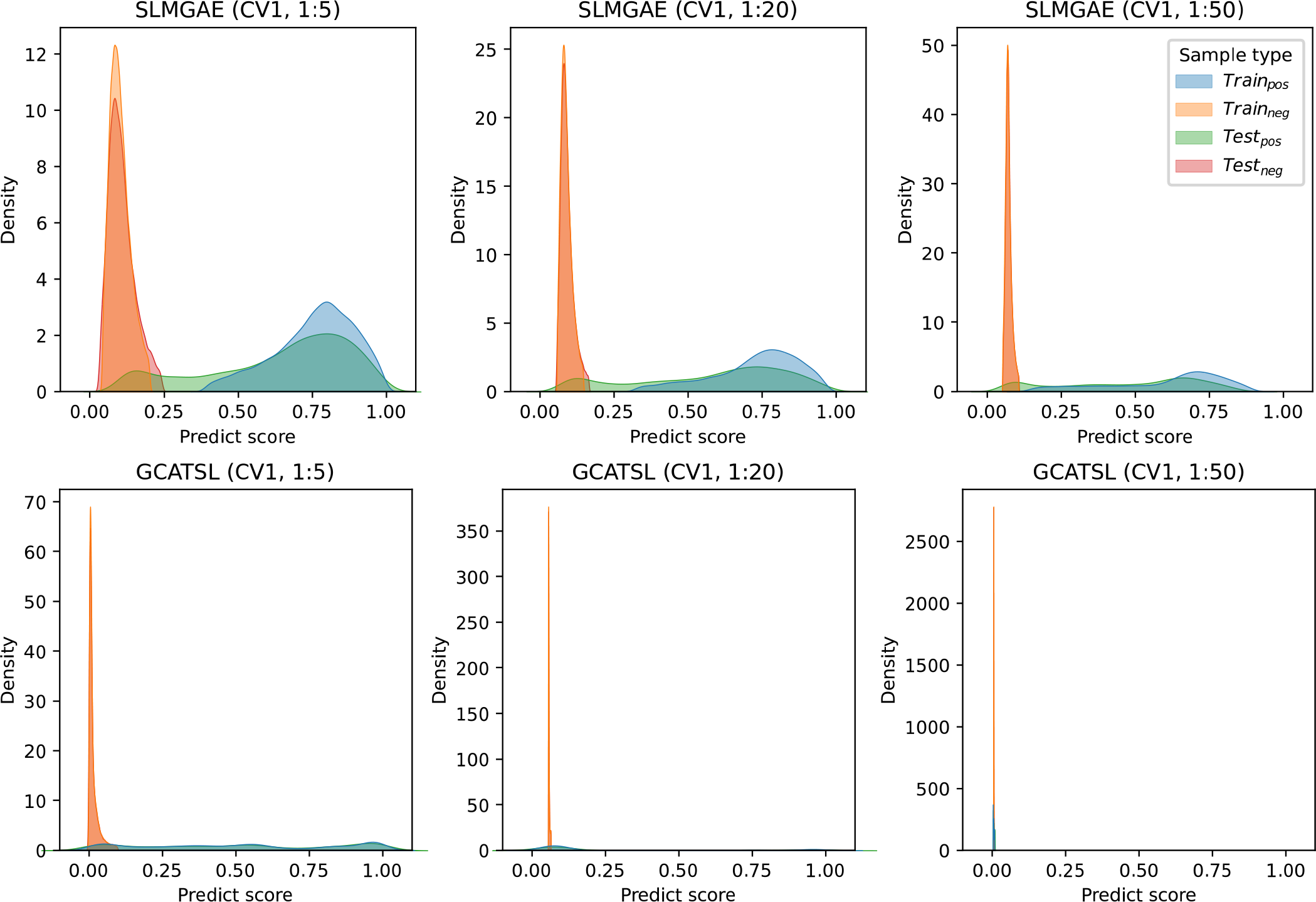
Distribution of predicted score. The score of gene pairs in the training and testing sets for the SLMGAE and GCATSL models are displayed in the top and bottom rows, respectively, for various positive and negative sample ratios (PNRs). The results are obtained with NSM_Rand_.

### Impact of negative sampling

Obtaining high-quality negative samples is crucial for the performance of the models. However, in the context of SL prediction, high-quality negative samples are scarce. Therefore, it is important to explore efficient and straightforward methods for obtaining high-quality negative samples from unknown samples. In this study, we evaluated three negative sampling approaches, namely NSM_Rand_, NSM_Exp_, and NSM_Dep_, which represent unconditional random negative sampling, negative sampling based on gene expression correlation, and negative sampling based on dependency score correlation, respectively (see Methods for details). Among these approaches, NSM_Rand_ has been widely used in existing SL prediction methods, and thus it will be used as the baseline for comparison. We denote the scenarios as (NSM_N_, CV1, 1:1), where N = Rand, Exp or Dep.

Based on the findings from the previous two subsections, certain characteristics regarding the model’s generalizability and robustness in the context of NSM_Rand_ can be observed. Here, we investigated two additional negative sampling methods (NSM_Exp_ and NSM_Dep_). Our observations revealed that models utilizing negative samples from NSM_Exp_ demonstrate improved classification performance compared to NSM_Rand_. On the other hand, models employing negative samples from NSM_Dep_ do not show significant performance differences relative to NSM_Rand_ (see Supplementary Data 1, 2, and 3). The results of the classification and ranking tasks are presented in Figure 5A and D. The majority of models demonstrate a marked improvement in the classification task when using NSM_Exp_, with GCATSL’s Classification score increasing from 0.709 to 0.808. Other models such as SLMGAE, GRSMF, KG4SL, DDGCN, and PiLSL all achieved Classification scores above 0.720. On the other hand, SL^2^MF and CMFW experienced a decrease in performance. In the ranking task, the performance of NSM_Dep_ was better than NSM_Exp_, but not as good as NSM_Rand_. CMFW, DDGCN, and SLMGAE experienced a considerable decrease in their Ranking scores when using either NSM_Dep_ or NSM_Exp_, while the other models did not demonstrate a significant variation.

**Figure 5.**
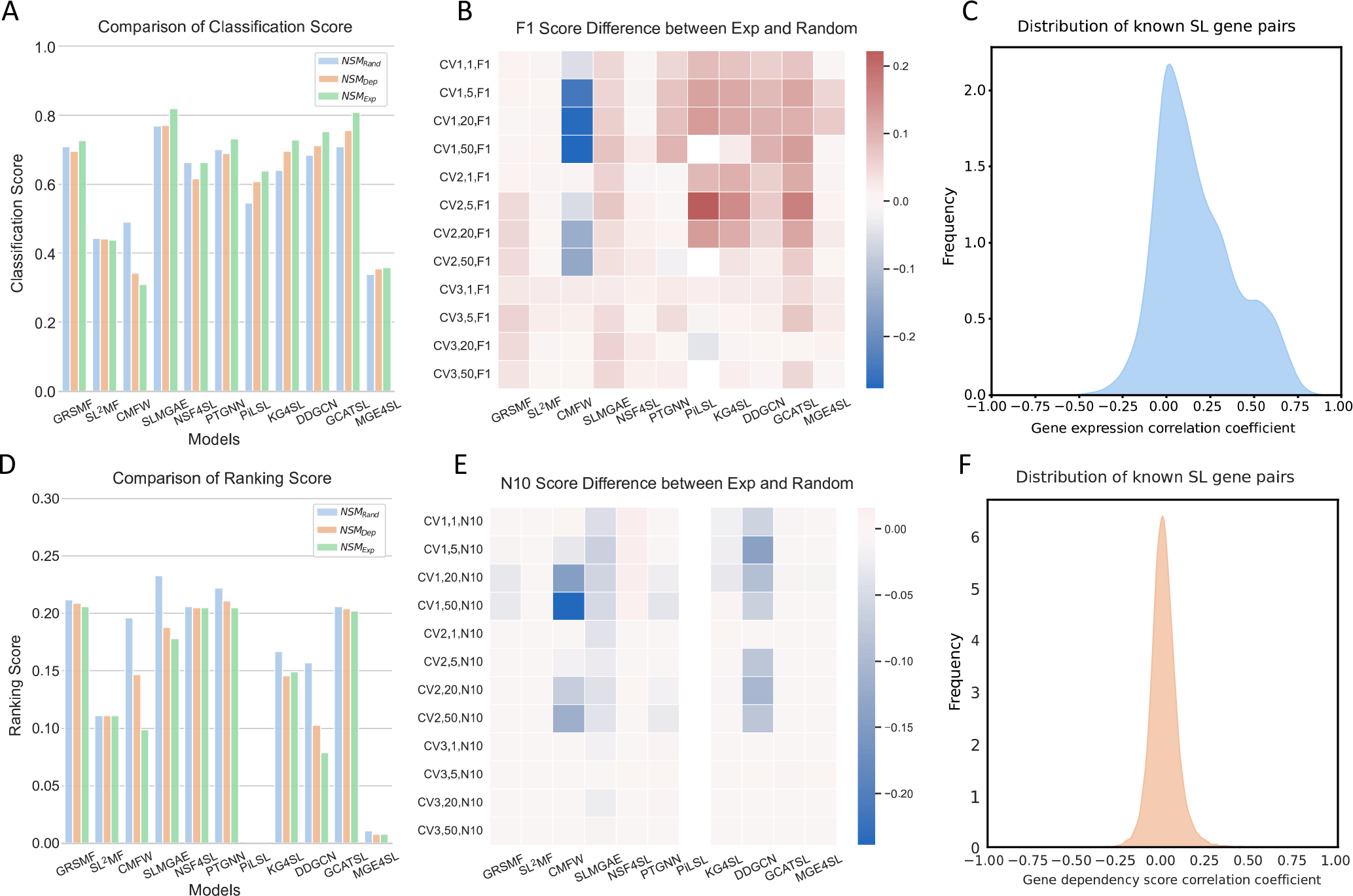
Comparative analysis of model performance and positive sampls. **A** and **D** show the overall score comparison of the model under classification and ranking tasks under all three negative sampling methods (NSMs). **B** and **E** illustrates the differences between classification and ranking tasks using NSM_Exp_ and NSM_Rand_ in different scenarios, as measured by the F1 score and N10 metrics. Red and blue signify an increase and a decrease, respectively, with darker shades indicating a larger difference. **C** and **F** denote the correlation coefficient distribution of known SL gene pairs under different data sources.

We also assessed the impact of negative sampling on the generalization and robustness of the model. As shown in Figure 5B and E, the negative samples obtained through NSM_Exp_ have a small impact on the performance of matrix factorization based methods, except for the CMFW model, which has a significant decrease in performance in classification tasks compared to NSM_Rand_. On the contrary, for deep learning based methods, the negative samples obtained through NSM_Exp_ improve the classification task performance of the model in various scenarios, especially for the CV1 and CV2 scenarios. It is noteworthy that the performance of the NSF4SL model is not affected by the quality of negative samples, as it “does not use negative samples” (i.e. does not use negative samples at all) during the training process. For ranking tasks, except for SLMGAE and DDGCN, the quality of negative sample has a relatively small impact on most models.

Furthermore, we investigated the potential reasons underlying the different impacts of the negative sampling method. As shown in Figure 5C, the distribution of gene expression correlation scores for known SL gene pairs exhibits a bias towards positive scores. Note that NSM_Exp_ selects gene pairs with negative correlation coefficient of gene expression. It is possible that, distinguishes the distribution of positive and negative samples in advance, reducing the difficulty of classification, it is able to because NSM_Exp_ improve the performance of the models in the classification task. By contrast, as shown in Figure 5F there is a symmetric distribution of correlation based on dependency scores, hampering the model’s ability to learn more effective features from the negative samples selected by NSM_Dep_.

## Discussion

Here, we present a comprehensive benchmarking study of 11 machine learning methods for predicting SL interactions. We constructed a dataset from multiple sources and evaluated all the methods on this dataset. Our results demonstrate that the predictive capabilities of these methods vary under different experimental settings. Specifically, among the matrix factorization-based methods, GRSMF exhibited superior stability and accuracy relative to the other two methods. For the deep learning-based methods, SLMGAE showed the most promising results. Our benchmark framework can further evaluate new models, making evaluations more consistent. There are multiple aspects worth discussing.

Our evaluation showed that despite using only SL data, DDGCN can achieve performance comparable to models that rely on multiple external data sources (e.g., PTGNN), possibly due to its unique design of dual-dropout. In addition, although both SLMGAE and MGE4SL incorporate multiple graph views and attention mechanisms, SLMGAE demonstrates for superior performance compared to MGE4SL. Further analysis of the two methods reveals that SLMGAE optimizes the model through three distinct objectives corresponding to SL, PPI, and GO, respectively. It distinguishes the main view from supporting views and reconstructs multiple graphs for different views via graph auto-encoders (GAEs). By contrast, MGE4SL optimizes the model by fusing all the information using a cross-entropy loss, which may introduce noise. The auto-encoder has been shown^43^ to reduce information loss caused by Laplacian smoothing^44^. This may be one of the reasons why the SLMGAE performs so well. KG4SL and GCATSL exhibit noticeable performance degradation when PNR=1:50, i.e., with an overwhelming proportion of negative samples, indicating the models’ inability to discriminate between sparse positive and abundant negative samples. Furthermore, models that combine different types of data, such as GO and PPI, usually yield better results than DDGCN. However, if inappropriate methods are used, such combinations can be counterproductive, as in the case of MGE4SL. We also observed that although KG4SL utilizes multiple relationships contained in the knowledge graph, its performance lags behind that of SLMGAE in both classification and ranking tasks. This suggests that an excess of supplementary information may not necessarily yield better results. As a valuable data resource, the best way to utilize the KG is still worth exploring.

Based on our study, we identified several issues and exploring in SL prediction that require attention. First, most models do not consider the realistic imbalanced proportions of SL and non-SL relationships. Theoretically, SL interactions constitute only a small fraction of all gene-gene interactions. As a result, there are much more non-SL gene pairs than SL gene pairs. The model must be able to handle highly imbalanced data in order to generate the desired predictions in real-world applications. Second, the CV1 scenario’s mainly focus is limited to a gene set with known SL relationships. Compared to CV1, the CV2 scenario requires the models to apply known patterns to other genes and identify novel SL interactions. That is, it evaluates a model’s ability to generalize known patterns to genes that are not yet involved in any known SL pairs. However, most methods currently do not take this situation into account. Third, the lack of high-quality negative samples can affect the evaluation of the model. Generally, when evaluating the performance of a discriminant model, it is necessary to test the model with both positive and negative samples of known labels. However, in the context of SL prediction, obtaining high-quality negative samples is often a daunting task. The false negative samples included in the test dataset could potentially skew the evaluation metrics. In recent years, with the development of CRISPR technology, there have been many works^45–47^using combinatorial CRISPR/Cas9^48^ to screen SL gene pairs, and the data generated from these experiments can provide reliable negative samples. But for a large space of gene pairs, the wet-lab method is still labor-intensive. Finally, it is essential to incorporate multiple data sources, such as GO and PPI, to enhance the generalizability of the model. Additional data related to genes and genetic interactions cover more extensive features of genes than the known SL gene set, these data can provide information for predicting new SL interactions. Additionally, KG is a valuable data source because it contains information on multiple aspects of a gene and is organized in a graph format that can be more interpretable. However, the current methods for predicting SL interactions still very rudimentary in harnessing KG’s potential.

There are a few possible future directions in the field of SL prediction by machine learning. Firstly, predicting SL interactions taking into account cancer-specific genetic context is key to developing precision cancer medicine. SL interactions are context-specific, meaning that the SL relationship between a pair of genes may only occur in one type of cancer (or cell line) but not in another. Wan et al. proposed EXP2SL^49^, a method that leverages unlabeled SL data to forecast cell line-specific SL pairs. The study showcased the effectiveness of L1000 expression profiles^50^ as feature data for SL prediction. Secondly, SL interactions can support drug repositioning. SL can help identify new targets of approved drugs or discover new combination treatment strategies. Zhang et al. developed SLKG^51^, which provides a computational platform for SL-based tumor therapy, and demonstrated that SLKG can help identify promising repurposed drugs and drug combinations. Thirdly, the concept of SL needs extension and refinement. Several classes of SL, such as synthetic dosage lethality (SDL)^52–54^ and collateral SL^55,56^, have been proposed by researchers to capture the inherent complexity of SL. Furthermore, Li et al.^57^ categorized SLs into two types: non-conditional/original and conditional SL. Conditional SL refers to the occurrence of synthetic lethal interactions that occur under specific conditions, such as genetic background, hypoxia, high levels of reactive oxygen species (ROS)^58^, and exposure to DNA damaging agents and radiation. The distinction between the various SL definitions is often overlooked in the current models. When these mixed data are input into the model, they may interfere with the model’s judgment of a specific type of SL. Fourthly, the lack of high-quality negative samples poses a challenge for researchers in SL prediction. While NSF4SL^42^ avoids this problem by using a contrastive learning framework to despense with the need of using negative samples, most classification models still rely on them. Lastly, most current machine learning models lack interpretability, making it difficult to assess the reliability of their predictions. Although performing well in terms of accurate and sensitive prediction, these models often do not incorporate underlying biological mechanisms. Improving the interpretability of the machine learning models can make them more practically useful and informative for users, including experimental biologists. Recently, Zhang et al. introduced KR4SL^59^, a novel SL prediction model based on KG reasoning. This model obtains the semantic representation of gene pairs by encoding the structural information of the directed subgraphs in KG and the textual semantics of entities, and further enhances the sequence (the path within a relational digraph) semantics with the help of gated recursive units (GRUs). KR4SL can provide clear explanations by uncovering the prediction process and providing evidence about the biological mechanisms of SL.

## Methods

### Data

For this benchmarking study, we obtained synthetic lethality (SL) label data from the database of SynLethDB 2.0, which contains a knowledge graph named SynLethKG^28^. In addition, we collected other data used by some models studied here including protein-protein interaction (PPI), gene ontology (GO), pathways, protein complexes, and protein sequences. For more details on the model data requirements, see Table 4.

To ensure consistency, we standardized the gene names in the data using the HUGO Gene Nomenclature Committee (HGNC)^60^ and the Ensembl Database^61^.

**Synthetic lethality labels**. After filtering, we finally obtained a dataset consisting of 9,845 unique genes and 35,913 positive SL gene pairs from the SynLethDB. We collected 54,012 entities and 2,233,172 edges from the SynLethKG. To facilitate integration, all the data was aligned by gene names. Next, we introduce the way to pre-process prior knowledge.

**Protein-protein interaction (PPI)**. We extracted protein interaction data from the BioGRID database^30^ to construct a PPI network, where nodes represent a subset of 9,845 genes while edges represents the interaction relationships between the nodes. Additionally, we reimplemented the FSweight algorithm^62^ on the PPI network to calculate the functional similarity matrix of proteins as input of SL^2^MF.

**Gene ontology (GO)**. We collected Gene Ontology (GO) data from the Gene Ontology database^29,63^, which consists of Gene Ontology Annotation (GOA) and GO terms. To compute the functional similarity between genes and the semantic similarity between GO terms, we employed the R package GOSemSim^64,65^.

**Pathway**. We retrieved pathway data from multiple databases, including KEGG^31^, Reactome^66^, and MSigDB^67,68^. In particular, we extracted the genes belonging to the same pathway and use them to construct a symmetrical mask matrix. In the matrix, a value of 1 is assigned when two genes are in the same pathway, and 0 otherwise.

**Dependency score and gene expression**. We collected gene expression and gene dependency score data from different cell lines in the DepMap^69^ database. Using these data, we designed two new negative sampling methods to evaluate the impact of negative samples on models performance.

### Experimental scenarios

#### Different proportions of negative samples

To evaluate the performance of synthetic lethality (SL) prediction methods, it is crucial to use realistic scenarios for training and testing. In most existing methods, the number of negative samples is set equal to the number of positive samples, i.e., a positive-negative ratio (PNR) of 1:1. However, in real data, there are much more non-SL gene pairs than SL gene pairs. Therefore, we tested four different PNRs in our experiments, i.e., 1:1, 1:5, 1:20, and 1:50.

#### Data split methods

Suppose *H* = {g_a_,g_b_, …g_n_} is the set of all human genes, and *P*_*SL*_ = {(g_i_,g_j_), (g_k_,g_l_), …} is the set of all known human SL gene pairs. The human gene set *H* can be divided into two sets, H_seen_ and H_unseen_, which represent currently known genes with SL interactions and the rest genes, respectively. Clearly *H*_*seen*_ ∪*H*_*unseen*_ = *H* and *H*_*seen*_ ∩*H*_*unseen*_ = /0. For the machine learning models in this benchmarking study, there are generally three situations encountered in the actual prediction of whether a pair of genes (g_a_, g_b_) have SL interaction: (1). {(g_a_,g_b_)|g_a_ ∈ *H*_*seen*,_ g_b_ ∈ *H*_*seen*_}, (2). {(g_a_,g_b_)|g_a_ ∈ *H*_*seen*,_ g_b_ ∈ *H*_*unseen*_}, and (3). {(g_a_,g_b_)|g_a_ ∈ *H*_*unseen*_, g_b_ ∈ *H*_*unseen*_}.

We simulated these possible scenarios through three data split methods (DSMs), namely CV1, CV2, and CV3 (Figure 6B).

**Figure 6.**
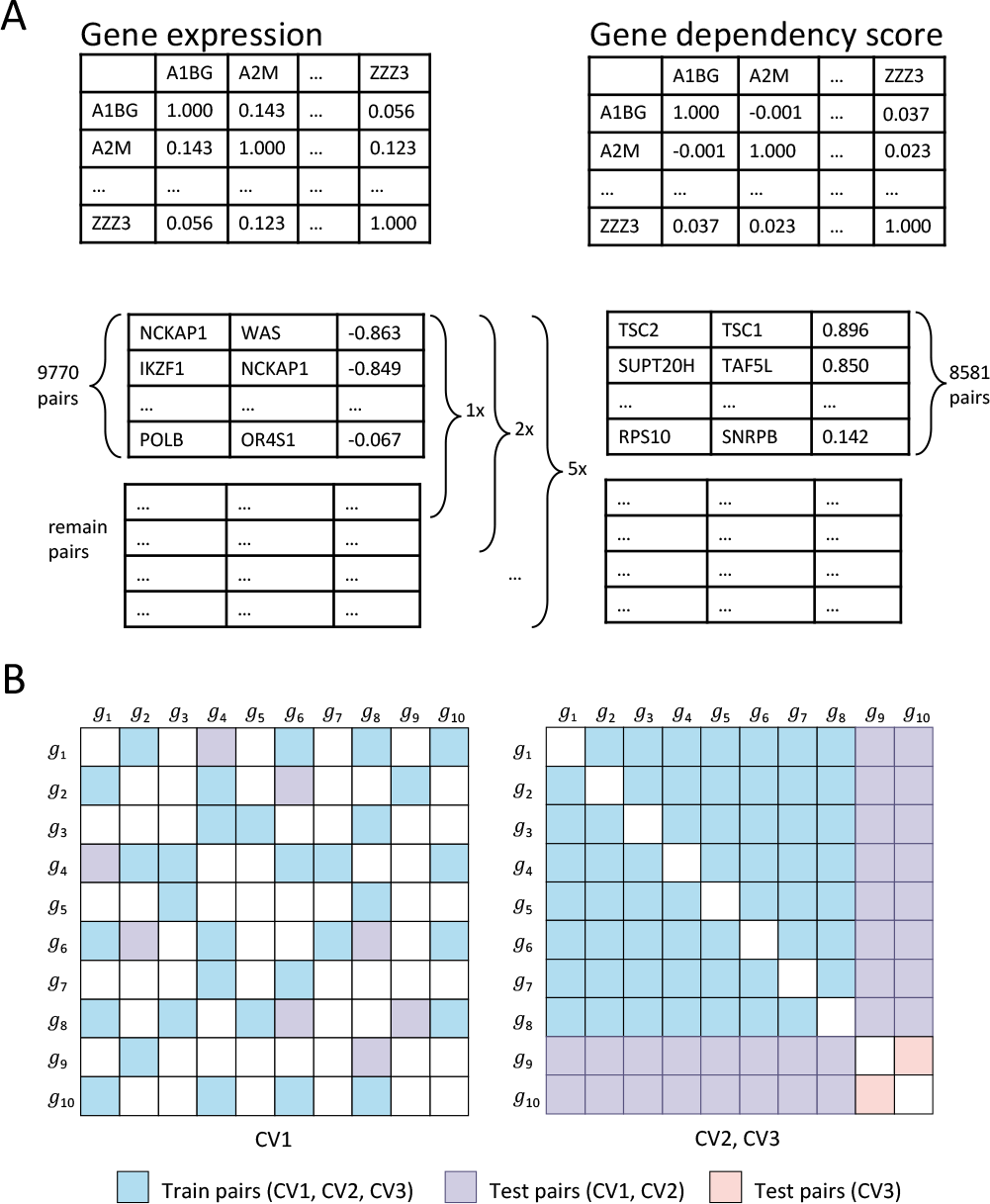
Schematic diagram of negative sampling strategy and data splitting method. **A** depicts the negative sampling steps based on gene expression and gene dependency scores. **B** illustrates the different data partitioning methods: the left matrix shows an example when DSM=CV1, with the blue and purple areas representing the randomly sampled training and test samples, respectively. For DSM=CV2 or CV3, their training samples are both drawn from the blue area, while the purple and orange represent the test sample regions for CV2 and CV3 respectively.

The settings are as follows:

1. CV1: we split the data into training and testing sets by SL pairs, where both genes of a tested pair may have occurred in some other gene pairs in the training set.
2. CV2: we split the data by genes, where only one gene of a tested pair is present in the training set.
3. CV3: we split the data by genes, where neither of a tested pair of genes is present in the training set.

#### Negative sampling methods

To train the deep learning models for SL prediction, a sufficient number of gene pairs are required, including negative samples. However, non-SL gene pairs are rarely known, which makes it difficult to satisfy the requirements of deep learning models. Therefore, negative sampling is often needed to obtain negative SL data for learning. A common strategy is to randomly select gene pairs from unknown samples as negative samples, which may include false negatives.

To address this issue, we designed two new negative sampling methods (NSMs) based on the DepMap database^69^. The DepMap database includes gene expression data, gene mutation data, gene dependency scores data, etc. of many cell lines. The gene dependency scores were assessed through the utilization of CRISPR technology, which involves the examination of cellular activity after the single knockout of a specific gene, and a higher gene dependency score indicates lower cell activity. From DepMap database, we obtained gene expression data and gene dependency scores obtained by CRISPR knockout experiments. We have designed two new negative sampling methods (NSMs) using these data: NSM_Exp_ and NSM_Dep_.

The NSM_Exp_ is based on the correlation of expression between two genes. For each pair of genes, we calculated the correlation coefficient of expression (corr_Exp_) between the genes across the cell lines. From Figure 5C, we observe that known SL gene pairs tend to have positive correlations of gene expression, i.e., corr_Exp_ > 0. Therefore, our sampling step is as follows:

1. We arranged all gene pairs in ascending order of their correlation scores;
2. To ensure that each gene appears in the negative samples, we first traverse the gene pairs sequentially from the beginning to find the smallest set of gene pairs that can contain all genes;
3. From the remaining samples, we extract them in order (ascending order in NSM_Exp_ and descending order in NSM_Dep_) and stop when the quantity reaches one, five, twenty, and fifty times the number of positive samples.

Similarly, NSM_Dep_ is based on the correlation of the gene dependency score. For each pair of genes, we calculated the correlation coefficient of the dependency score (corr_Dep_) between the two genes. We found that the corr_Dep_ of the pairs of known SL genes were distributed mainly in the range [*−* 0.2, 0.2]. Therefore, we first take absolute values for all corr_Dep_ and use a sampling step similar to NSM_Exp_, but unlike NSM_Exp_, in the first step of the sampling, we rank all gene pairs according to their correlation scores from the highest to the lowest (Figure 6A shows the specific negative sampling steps.).

### Evaluation metrics

To comprehensively evaluate the performance of the models, we utilized six metrics. For the classification task, we used three metrics: area under the receiver operating characteristic curve (AUROC), area under the precision-recall curve (AUPR), and F1 score. These metrics are commonly used for binary classification. For the gene ranking task, we employed three metrics: normalized discounted cumulative gain (NDCG@K), Recall@K, and Precision@K. NDCG@K measures whether the known SL gene pairs are in a higher position in the predicted list of a model, while Recall@K and Precision@K are used to evaluate the model’s ability to measure its coverage of relevant content and accuracy in returning the top-K results, respectively. The definitions of these metrics are as follows:

- Area Under the Receiver Operating Characteristic Curve (AUROC): AUROC measures the model’s ability to classify samples at different thresholds. It is calculated as the area under the receiver operating characteristic curve, which is a curve plotted with false positive rate on the x-axis and true positive rate on the y-axis. The value of AUROC ranges between 0 and 1.
- Area Under the Precision-Recall Curve (AUPR): AUPR is a performance metric used to evaluate binary classifiers, which measures the average precision across different recall levels. Like AUROC, AUPR can be used to evaluate the performance of classifiers in the presence of imbalanced classes or uneven sample distributions. AUPR is a more sensitive metric than AUROC, particularly for classification of imbalanced data.
- F1 score: The F1 score is a metric for evaluating the overall effectiveness of a binary classification model by considering both precision and recall. Combining precision and recall into a single value, it provides a balanced measure of the effectiveness of the model.
- Normalized discounted cumulative gain (NDCG@K): NDCG@K can be used to evaluate the ability of a model in ranking candidate SL partners for a gene *g*_*i*_. NDCG@K is calculated as NDCG@K = DCG@K/IDCG@K, where IDCG@K is the maximum DCG@K value among the top-K predictions, and DCG@K is calculated as:

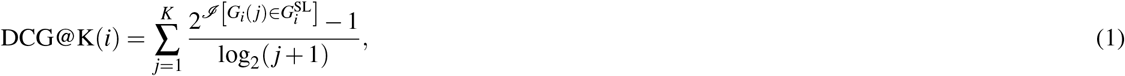

where 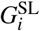 denotes all known genes that have SL relationships with gene *g*_*i*_, *G*_*i*_(*j*) is the *j*-th gene on the list of predicted SL partners for gene *ℐ*_*i*_, and *I* [*·*] is the indicator function.
- Recall@K: Recall@K measures the proportion of correctly identified hits among the top K predicted SL partners to the total known SL partners for gene *g*_*i*_.

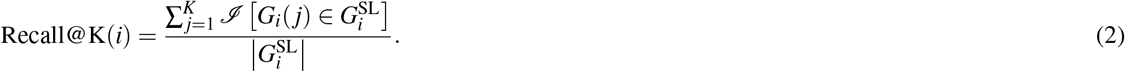
- Precision@K: Precision@K represents the proportion of correctly identified SL partners among the top K predicted SL partners of gene *g*_*i*_.

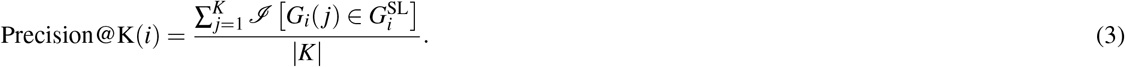
- To evaluate the overall performance of a model, we calculate its performance under classification and ranking tasks separately, and combine them with equal weights to obtain an indicator that reflects the overall performance of the model, i.e., Overall = (Classification score + Ranking score) / 2. The classification and ranking scores are calculated as follows, respectively:

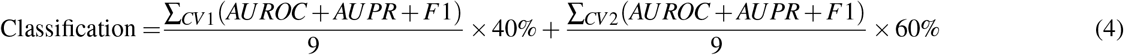

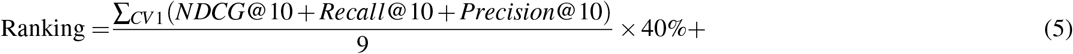

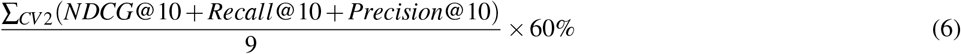

In the context of predicting new SL relationships, the CV2 scenario is more realistic and prevalent, and thus it holds greater significance. For most models, CV3 is overly challenging. Therefore, when calculating the integrative classification and ranking scores, we do not include scores under CV3 and set the weights for CV1 and CV2 scenarios to 40% and 60% respectively.

### Model selection and implementation

In this study, We benchmarked 11 *in silicon* methods for synthetic lethality prediction, including 3 matrix factorization-based methods and 8 deep learning-based methods (Table 3). The details of these methods can be found in the Supplementary 1 and 2. For all the methods, their implementation details are as follows:

GRSMF^34^: due to the lack of executable code in the code repository of the method itself, the code version we used is GRSMF implemented in GCATSL https://github.com/lichenbiostat/GCATSL/tree/master/baseline%20methods/GRSMF. We set *num_nodes* to 9845, which is the number of genes in our data.

SL^2^MF^32^: we used the code of SL^2^MF from https://github.com/stephenliu0423/SL$^2$MF. The *num_nodes* was set to 9845.

CMFW^33^: we used the code of CMFW from https://github.com/lianyh/CMF-W.

SLMGAE^37^: we used the code of SLMGAE from https://github.com/DiNg1011/SLMGAE. We used the default settings.

NSF4SL^42^: we used the code of NSF4SL from https://github.com/JieZheng-ShanghaiTech/NSF4SL. The settings *aug_ratio* = 0.1 and *train_ratio* = 1 were used.

PTGNN^39^: we used the code of PTGNN from https://github.com/longyahui/PT-GNN. We have limited the maximum length of protein sequences to 600 and redesigned the word dictionary based on the original paper.

PiLSL^41^: we used the code of PiLSL from https://github.com/JieZheng-ShanghaiTech/PiLSL. We set the following parameters: –hop 3, –batch_size 512. When calculating the metrics for the ranking task, we need to calculate the scores of all gene pairs, which are about 50 million. PiLSL is a pair by pair prediction approach that demands significant time to obtain all the necessary scores. Therefore, we only considered the performance of the model under the classification task when the PNRs of 1:1, 1:5, and 1:20.

KG4SL^40^: we used the code of KG4SL from https://github.com/JieZheng-ShanghaiTech/KG4SL. The default parameters are used for the experiment.

DDGCN^35^: we used the code of DDGCN from https://github.com/CXX1113/Dual-DropoutGCN. We set dropout = 0.5 and lr = 0.01, which are the default parameter settings.

GCATSL^36^: we used the code of GCATSL from https://github.com/lichenbiostat/GCATSL. MGE4SL^38^: we used the code of DDGCN from https://github.com/JieZheng-ShanghaiTech/MGE4SL.

### Computational resource

The experiments were conducted on a workstation equipped with 4 Intel(R) Xeon(R) Gold 6242 CPUs @ 2.80GHz, with a total of 64 cores and 22,528 KB cache, along with 503 GB of memory. The system was also equipped with three Tesla V100s 32 GB GPUs, providing a total of 96 GB of GPU memory. The operating system used was Linux Ubuntu 20.04.

## Data availability

All the data used in our study include: SL labels from SynLethDB 2.0 https://synlethdb.sist.shanghaitech.edu.cn/#/download. The entrez IDs of the genes come from the NCBI database https://www.ncbi.nlm.nih.gov/gene/. Ensemble ID from https://asia.ensembl.org/index.html. The PPI data come from the data released by BioGRID on June 25, 2022, and the download link is https://downloads.thebiogrid.org/File/BioGRID/Release-Archive/BIOGRID-4.4.211/BIOGRID-ALL-4.4.211.tab.zip. GO annotation and GO term data are respectively from http://geneontology.org/gene-associations/goa_human.gaf.gz and http://geneontology.org/docs/download-ontology/#go_obo_and_owl. The gene expression data and gene dependency score data of the cell lines are from DepMap Public 22Q4 https://figshare.com/articles/dataset/DepMap_22Q4_Public/21637199/2. Pathway data are from KEGG database https://www.kegg.jp/kegg-bin/download_htext?htext=hsa00001&format=json&filedir=kegg/brite/hsa. The protein sequence data are from the UniProt database, released on July 22, 2022 https://www.uniprot.org/help/downloads.

## Code availability

The Python codes used for our benchmark are available at https://github.com/JieZheng-ShanghaiTech/SL_benchmark

## Author contributions statement

J. Z., M.W., and Y.F. conceived this idea, J.Z. and M.W. designed and guided the project, Y.F., Y.L., Q.L., M.W., and J.Z. participated in manuscript writing, Y.F. completed experiments and result analysis, Y.F. designed charts, H.W., and Y.O. assisted in literature review.

## Additional information

To include, in this order: **Accession codes** (where applicable); **Competing interests** (mandatory statement).

The corresponding author is responsible for submitting a competing interests statement on behalf of all authors of the paper. This statement must be included in the submitted article file.

