## Supplementary Materials for "Benchmarking Machine Learning Methods for Synthetic Lethality Prediction in Cancer"

### Artificial Intelligence for Synthetic Lethality Prediction in Cancer: A Survey of Machine Learning Methods and Evaluation

#### 1 Matrix factorization-based methods

### 1.1 SL$^2$MF

Liu et al. proposed the SL$^2$MF[1] model for predicting SL interactions, which is based on matrix factorization. Specifically, this model employs logistic matrix factorization (LMF) to map genes into a low-dimensional latent space and model the probabilities of SL interactions between genes using their latent representations. The matrix $\mathbf{U} \in \mathbb{R}^{n\times d}$ represents the latent vectors of each gene $\mathbf{U}_i \in \mathbb{R}^{1\times d}$, where $d$ is the dimensionality of the latent space. The probability of an SL interaction between two genes $g_i$ and $g_j$ is calculated using the following logistic function:

$$p_{ij} = \frac{1}{1+\exp\left(-\mathbf{U}_i\mathbf{U}_j^T\right)} \tag{1}$$

In order to improve the predictive accuracy of the model, the authors integrated knowledge from the PPI network and GO annotation into their approach. The authors hypothesized that genes with similar functional and/or network properties should exhibit similar representations in the latent space. To incorporate this information, they utilized additional loss functions in the model that take into account the GO and PPI similarities between genes for SL interaction prediction.

$$L_{\mathscr{G}} = \frac{1}{2}\sum_{i=1}^{n}\sum_{u_j\in N^G(u_i)} s_{ij}^G \left\|\mathbf{U}_i - \mathbf{U}_j\right\|_2^2 \tag{2}$$

$$L_{\mathscr{P}} = \frac{1}{2}\sum_{i=1}^{n}\sum_{u_j\in N^P(u_i)} s_{ij}^P \left\|\mathbf{U}_i - \mathbf{U}_j\right\|_2^2 \tag{3}$$

where $N^G$ and $N^P$ represent the nearest neighbors of each gene in the GO and PPI similarity matrices, respectively.

##### 1.2 GRSMF

Huang et al. proposed a graph regularized self-representative matrix factorization (GRSMF) model[2], which is based on the self-representative matrix factorization (SMF)[3] method. In traditional SMF models, the input data $\mathbf{X}$ is self-represented by a linear combination of its columns as $\mathbf{X} \approx \mathbf{X}\mathbf{U}$, where $\mathbf{U} \in \mathbb{R}^{n\times n}$ is the coefficient matrix that represents the columns of $\mathbf{X}$. Since the SL interaction matrix is symmetric, the representation of its rows should be the same as the representations of its columns, and thus the self-representation of the symmetric matrix $\mathbf{X}$ is given by $\mathbf{X} \approx \mathbf{U}^T\mathbf{X}\mathbf{U}$. For the $i$th gene $g_i$, $\mathbf{U}_{li}$ denotes the probability of gene $g_i$ being represented by $g_l$, which captures the similarity between $g_i$ and $g_l$ based on their SL interactions with other genes. By taking into account some constraints, the following regularized self-representative matrix factorization model can be built

$$\min_{\mathbf{U}} \left\|\mathbf{X} - \mathbf{U}^T\mathbf{X}\mathbf{U}\right\|_F^2 + \lambda\left\|\mathbf{U}\right\|_F^2 \tag{4}$$

$$s.t. \quad 0 \le \mathbf{U} \le 1, \quad \sum_{l=1}^{n}\mathbf{U}_{li} = 1, \quad for \quad i = 1,\dots,n, \tag{5}$$

where $\|\cdot\|_F$ is the Frobenius norm and $\lambda$ is a tuning parameter which controls the influence of the $l_2$ regularization.

Due to the limited number of known SL interactions, it can be challenging to learn a comprehensive representation matrix. To address this limitation, the authors incorporated prior information that captures similarities among genes, by introducing a graph regularization term to the SMF model. Specifically, let $\mathbf{S} \in \mathbb{R}^{n\times n}$ denote the GO similarity matrix, where $\mathbf{S}_{ij}$ represents the functional similarity between genes $g_i$ and $g_j$. The graph regularization based on $\mathbf{S}$ is defined as follows:

$$R = \frac{1}{2}\sum_{l=1}^{n}\sum_{i,j=1}^{n}\left\|\mathbf{U}_{il} - \mathbf{U}_{jl}\right\|^2\mathbf{S}_{ij} = \mathrm{Tr}\left(\mathbf{U}^T\mathbf{L}\mathbf{U}\right) \tag{6}$$

where $\mathrm{Tr}(\cdot)$ denotes the trace of a matrix, $\mathbf{D}$ is a diagonal matrix with $d_{ii} = \sum_{j=1,n} \mathbf{S}_{ij}$ and $\mathbf{L} = \mathbf{D} - \mathbf{S}$.

Above all, the final objective optimization function of GRSMF is formulated as

$$\min_{\mathbf{U}} \left\|\mathbf{X} - \mathbf{U}^T\mathbf{X}\mathbf{U}\right\|_F^2 + \lambda \|\mathbf{U}\|_F^2 + \beta\, \mathrm{Tr}\left(\mathbf{U}^T\mathbf{L}\mathbf{U}\right) \tag{7}$$

$$s.t. \quad 0 \leq \mathbf{U} \leq 1, \quad \sum_{l=1}^{n} \mathbf{U}_{li} = 1, \quad for \quad i = 1, \ldots, n. \tag{8}$$

where $\beta$ controls the effect of graph regularization.

##### 1.3 CMF-W

Biological data sets are usually represented as matrices that include paired relationship data between two entities. Matrix sets can have many relationships between organisms. Collective matrix factorization (CMF)[4] and its extension are models designed to collectively learn numerous such connections. Liany et al. proposed a new method called CMF-W[5], which is an extension of the CMF approach. Specifically, the authors introduced three improvement measures:

First, generate the feature vector of the matrix using principal component analysis (PCA). When performing dimensionality reduction using PCA, the authors selected the minimum number of principal components required to achieve a cumulative explained variance ratio greater than 0.9 as the dimension of the reduced matrix. Second, provide the graph-based features that are matrix-specific. Including node degree, compactness centrality, intermediate centrality[6], information centrality[7], feature vector centrality[8], Gil Schmidt power index[9] and Flow Between score[10]. Third, incorporating the matrix-specific weight matrix.

In the CMF-W method, each matrix is modeled as the product of three factors:

$$\mathbf{X}^{(m)} \approx \mathbf{U}^{(r_m)}\mathbf{U}^{(c_m)^T}\mathbf{W}^{(m)} \tag{9}$$

The first two items are consistent with the original CMF method, that is, the row and column entity representation of the matrix, and the third item is the matrix specific weight matrix $\mathbf{W}^{(m)}$. For two matrices with the same row and column potential factors, $\mathbf{W}^{(m)}$ can be different.

The latent factors (including $\mathbf{W}^{(m)}$) are learned by solving the optimization problem:

$$\min_{\mathbf{U},\mathbf{W}} \sum_{m=1}^{M} d\left(\mathbf{X}^{(m)}, \mathbf{U}^{(r_m)}\mathbf{U}^{(c_m)^T}\mathbf{W}^{(m)}\right) \tag{10}$$

where $d$ is the Frobenius norm of the difference between $\mathbf{X}^{(m)}$ and $\mathbf{U}^{(r_m)}\mathbf{U}^{(c_m)^T}\mathbf{W}^{(m)}$. For $m \times n$ matrix $\mathbf{X}^{(m)}$ and latent dimension $k$, the dimensions of $\mathbf{U}^{(r_m)}$, $\mathbf{U}^{(c_m)^T}$, $\mathbf{W}^{(m)}$ are $m \times k$, $n \times k$, $n \times n$, respectively.

CMF-W overcomes the issue that arises when several input matrices include the same entity type, the classic CMF cannot learn the unique representation of each entity. Furthermore, the revised model can be directly utilized for input data and derived features, minimizing the effort of feature engineering.

#### 2 Graph neural network methods

##### 2.1 DDGCN

Due to the extreme sparsity of known SL interactions, overfitting is a common issue when training on such a sparse graph. While dropout is usually used to prevent overfitting, the effect of traditional dropout may be inadequate for SL interaction prediction. To enhance the robustness of gene representation for SL prediction, Cai et al. proposed a more effective and robust double dropout mechanism based on GCN, called DDGCN[11]. This model is composed of two paths with shared parameters that model the dual forms of dropout, namely coarse-grained node dropout and fine-grained edge dropout. The DDGCN model is designed to improve the accuracy of SL interaction prediction by enabling more robust gene representation.

**Coarse-grained node dropout** is achieved by using the identity matrix $\mathbf{I}$. Dropout is applied to the identity matrix, and subsequently encode the first-layer representation of each gene using graph convolution:

$$\mathbf{H}_{\mathbf{I}}^{(1)} = \mathrm{ReLU}\left(\hat{\mathbf{A}}\left(\mathbf{M}_{\mathbf{I}}^{(1)} \odot \mathbf{I}\right)\mathbf{W}^{(1)}\right), \tag{11}$$

where $\odot$ is the element-wise multiplication. $\mathbf{M}_{\mathbf{I}}^{(1)} \in \mathbb{R}^{n\times n}$ is the dropout mask in the first layer. $\mathbf{W}^{(1)} \in \mathbb{R}^{n\times d_1}$ is a trainable weight matrix for the first layer. $\hat{\mathbf{A}} = \mathbf{D}^{-\frac{1}{2}}\tilde{\mathbf{A}}\mathbf{D}^{-\frac{1}{2}}$ is the normalized graph adjacency matrix following Kipt et al.[12], where $\tilde{\mathbf{A}} = \mathbf{A} + \mathbf{I}$, $\mathbf{A}$ is the adjacency matrix and $\mathbf{D} \in \mathbb{R}^{n\times n}$ is a diagonal matrix which the diagonal elements are defined as $d_{ii} = \sum_{j=1}^{n} \tilde{\mathbf{A}}_{ij}$.

**Fine-grained edge dropout** is achieved by using the adjacency matrix $\mathbf{A}$, where the $i$th row or column of $\mathbf{A}$ corresponds to the SL interactions between gene $g_i$ and others. Authors apply dropout to $\mathbf{A}$ as follows

$$\mathbf{H}_{\mathbf{A}}^{(1)} = \mathrm{ReLU}\left(\hat{\mathbf{A}}\left(\mathbf{M}_{\mathbf{A}}^{(1)} \odot \mathbf{A}\right)\mathbf{W}^{(1)}\right) \tag{12}$$

$$\mathbf{H}_{\mathbf{A}}^{(2)} = \hat{\mathbf{A}}\left(\mathbf{M}_{\mathbf{A}}^{(2)} \odot \mathbf{H}_{\mathbf{A}}^{(1)}\right)\mathbf{W}^{(2)} \tag{13}$$

where $\mathbf{M}_{\mathbf{A}}^{(l)}$ is the dropout mask in the $l$th layer.

An important aspect of this model is parameter sharing, which allows the GCN parameters $\mathbf{M}^{(l)}$ in each layer to be shared by the dual dropouts.

The DDGCN model generates two embedding matrices $\mathbf{H}_{\mathbf{I}}^{(2)}$ and $\mathbf{H}_{\mathbf{A}}^{(2)}$ through the two paths. To estimate the interaction confidence of each gene pair $(g_i, g_j)$, the following formula will be used:

$$\hat{y}_{\mathbf{I}}(i,j) = \mathrm{Dec}\left(\mathbf{H}_{\mathbf{I}}^{(2)}\right) \tag{14}$$

$$\hat{y}_{\mathbf{A}}(i,j) = \mathrm{Dec}\left(\mathbf{H}_{\mathbf{A}}^{(2)}\right) \tag{15}$$

where $\mathrm{Dec}(\cdot)$ is the inner-product decoder[13] defined as

$$\mathrm{Dec}(\mathbf{U}) = \sigma\left(\mathbf{U}\mathbf{U}^{\top}\right) = \frac{1}{1+\exp\left(-\mathbf{U}\mathbf{U}^{\top}\right)} \tag{16}$$

Let $y_{ij} = 1$, if $(g_i, g_j)$ is a known SL interaction, otherwise $y_{ij} = 0$. Thus, the overall loss function can be formulated as follows:

$$L = \sum_{i=1}^{n}\sum_{j=i+1}^{n} \mathrm{CE}\left(y_{ij}, \hat{y}_{\mathbf{I}}(i,j)\right) + \alpha\,\mathrm{CE}\left(y_{ij}, \hat{y}_{\mathbf{A}}(i,j)\right), \tag{17}$$

where CE is the cross entropy loss, $\alpha > 0$ is a hyperparameter controlling the trade-off between the two paths. For the final prediction, they use the geometric mean to aggregate the two prediction scores $\hat{y}_{\mathbf{I}}(i,j)$ and $\hat{y}_{\mathbf{A}}(i,j)$:

$$\hat{y}(i,j) = \sqrt[1+\alpha]{\hat{y}_{\mathbf{I}}(i,j) \times \hat{y}_{\mathbf{A}}(i,j)^{\alpha}} \tag{18}$$

#### 2.2 MGE4SL

In order to tackle the sparsity of SL data, Lai et al.[14] suggested a multi graph integrated network structure, and added seven additional biological features by integrating other gene relational databases (such as GO, Corum, Reactome, etc.) to improve the performance of SL prediction.

For each graph, a two-layer GCN is used to generate the embeddings of the nodes. The layers are formulated as follows:

$$\mathbf{H}_{\mathbf{A}}^{(1)} = \mathrm{ReLU}\left(\hat{\mathbf{A}}\mathbf{X}\mathbf{W}^{(1)} + \mathbf{B}\right) \tag{19}$$

$$\mathbf{H}_{\mathbf{A}}^{(2)} = \hat{\mathbf{A}}\mathbf{H}_{\mathbf{A}}^{(1)}\mathbf{W}^{(2)} + \mathbf{B} \tag{20}$$

where the $\hat{\mathbf{A}}$ is defined in the same way as in the previous DDGCN.

Since the data from Corum were too sparse, the authors eventually used the other six graphs to obtain the multi-type node embeddings separately, and then they separately tried summing, splicing, and using CNNs to fuse these features and use them to predict SL interactions. Among these, concat method has better results. The author believes that this is because concat method keeps more knowledge information, and a simpler fully connected network is more successful than a complex CNN network.

##### 2.3 GCATSL

Long et al. proposed GCATSL[15], which is a graph context attention network based on GAT[16] that utilizes multiple biological data sources to generate diverse feature maps as input. The authors introduced a dual attention mechanism, operating at both the node and feature level, to capture the influence of local and global neighbors for learning gene expression from different feature maps. The extracted features were aggregated with the original features using a multilayer perceptron.

**Local representations** are obtained from the set of nodes that are directly connected to gene $g_i$, which are defined as its local neighbors. The authors introduce a node-level attention mechanism to learn the different levels of importance of nodes in local representations. Specifically, for a given gene $g_i$, the importance of its local neighbors is calculated as follows:

$$e_{ij}^{l}(g_i, g_j) = f(\mathbf{W}_1 h_i, \mathbf{W}_1 b_j), \tag{21}$$

where $e_{ij}^{l}$ is the attention score that indicates the importance of the neighbor $g_j$ to $g_i$. $f(\cdot)$ denotes a single-layer feed-forward neural network and $\mathbf{W}_1 \in \mathbb{R}^{d_1 \times d_2}$ is a learnable weight matrix. And $h_i$ is come from the feature matrix $\mathbf{H} \in \mathbb{R}^{n \times d_1}$, which is the GO or PPI similarity score matrix obtained by dimensionality reduction using PCA.

In order to make the attention scores comparable across different nodes, a softmax function is applied to normalize the scores across all local neighbors (note as $\mathcal{N}_i^{l}$) of gene $g_i$. The resulting normalized score is denoted as $\alpha_{ij}^{l}$.

Then, a new local representation $h_i^{l}$ for $g_i$ was derived by aggregating the representations of its local neighbors according to their attention coefficients. However, the attention of an individual node may be unstable and may introduce noise in the model. To address this issue and reduce the noise, the node attention is extended to multi-head attention by repeating the node attention mechanism $K$ times, and then concatenate the $K$ learned representations into gene $g_i$'s local representation as follows:

$$h_i^{l} = \|_{k=1}^{K} \operatorname{ReLU}\left(\sum_{j \in \mathcal{N}_i^{l}} \alpha_{ij}^{l} \cdot \mathbf{W}_1^{k} b_j\right) \tag{22}$$

where $\|$ denotes the operation of vector concatenation.

**Global representations** are obtained from the global neighbors of gene $g_i$. Global neighbors are defined as nodes that are at least two hops away from a given node in the graph. Intuitively, global neighbors can provide valuable information for modeling the center node. To this end, a random walk with restart (RWR)[17] based attention mechanism was employed to learn the node representations from the global neighbors.

After obtaining the global neighborhood of a node, similar to the local representations, the global representations of a node $h_i^{g}$ can be obtained by the following equation:

$$h_i^{g} = \|_{k=1}^{K} \operatorname{ReLU}\left(\sum_{j \in \mathcal{N}_i^{g}} \alpha_{ij}^{g} \cdot \mathbf{W}_1^{k} b_j\right) \tag{23}$$

To combine the representations more accurately, the bi-interaction aggregator is used, which encodes feature interactions through two MLPs. The bi-interaction aggregator is formulated as follows:

$$\begin{aligned} z_i = {} & \operatorname{LeakyReLU}\left(\mathbf{W}_2\left(h_i + h_i^{l} + h_i^{g}\right) + b_2\right) \\ & + \operatorname{LeakyReLU}\left(\mathbf{W}_3\left(b_i \left\| h_i^{l} \right\| h_i^{g}\right) + b_3\right), \end{aligned} \tag{24}$$

where $\mathbf{W}_2 \in \mathbb{R}^{Kd_2 \times d_3}$, $\mathbf{W}_3 \in \mathbb{R}^{3Kd_2 \times d_3}$, $b_2 \in \mathbb{R}^{d_3}$ and $b_3 \in \mathbb{R}^{d_3}$ are trainable weight operation. $z_i$ is the feature-specific representation of gene $g_i$.

As different feature graphs contain distinct contextual information, a feature-level attention was implemented to aggregate feature-specific representations. The attention score $w_i^{k}$ represents the importance of the $k$th feature graph to the gene $g_i$.

The final representation $\mathbf{Z}_i$ for $g_i$ is:

$$\mathbf{Z}_i = \sum_{k=1}^{T} \beta_i^k \cdot z_i^k \tag{25}$$

where $T$ is the number of feature graphs, $\beta_i^k$ is the normalized attention score using the softmax function. $z_i^k$ is the feature-specific representation of $g_i$.

The final probability score matrix $\mathbf{Q}$ and reconstruction loss are:

$$\mathbf{Q} = \text{sigmoid}\left(\mathbf{Z}\mathbf{W}_{de}\mathbf{Z}^T\right) \tag{26}$$

$$L_{Total} = \sum_{(i,j)\in\Omega^+\cup\Omega^-} \Phi\left(\mathbf{Q}_{ij}, \mathbf{A}_{ij}\right) + \gamma \parallel \mathbf{W}_{de} \parallel_F^2 \tag{27}$$

where $\mathbf{W}_{de}$ is a learnable latent factor that projects representations back to original feature space for genes. $\Phi(\cdot)$ is the MSE loss. $\gamma$ is weight factor that is used to control the impact of $\mathbf{W}_{de}$.

##### 2.4 SLMGAE

Hao et al. proposed a method for predicting SL interactions using Multi-view Graph Auto-Encoder, named SLMGAE[18]. In this approach, the authors utilized graphs from diverse data sources(e.g. PPI, GO, etc.) as support views, and the SL graph as the main view. They applied the GAE to reconstruct the graphs for different views and employed a two-layer GCN to calculate the node embedding matrix $\mathbf{Z}_2^m$ and reconstruct graph $\mathbf{S}^m$ for the main view using the following equations:

$$\mathbf{Z}_1^m = \text{LeakyReLU}\left(\hat{\mathbf{A}}\mathbf{F}\mathbf{W}_1^m\right) \tag{28}$$

$$\mathbf{Z}_2^m = \text{LeakyReLU}\left(\hat{\mathbf{A}}\mathbf{Z}_1^m\mathbf{W}_2^m\right) \tag{29}$$

$$\mathbf{S}^m = \mathbf{Z}_2^m\mathbf{W}_d^m\mathbf{Z}_2^{m\mathbf{T}} \tag{30}$$

where the $\hat{\mathbf{A}}$ is defined in the same way as in the previous DDGCN. $\mathbf{F}$ is the initial feature matrix. $\mathbf{W}_1^m$ and $\mathbf{W}_2^m$ are the weight matrices, and $\sigma_1$ and $\sigma_2$ are the activation functions in the first and second layers. $\mathbf{W}_d^m$ is a view specific trainable matrix in the decoder.

Similarly, for each support view $u$, the node embedding matrix $\mathbf{Z}_2^u$ and reconstruct graph $\mathbf{S}^u$ can be computed by

$$\mathbf{Z}_1^u = \text{LeakyReLU}\left(\hat{\mathbf{A}}^u\mathbf{F}\mathbf{W}_1^u\right) \tag{31}$$

$$\mathbf{Z}_2^u = \text{LeakyReLU}\left(\hat{\mathbf{A}}^u\mathbf{Z}_1^u\mathbf{W}_2^u\right) \tag{32}$$

$$\mathbf{S}^u = \mathbf{Z}_2^u\mathbf{W}_d^u\mathbf{Z}_2^{u\mathbf{T}} \tag{33}$$

Then the reconstruction loss $\mathscr{L}_M$ for the main view and $\mathscr{L}_S$ for the support views can be derived as follows

$$\mathscr{L}_M = \Phi\left(\mathbf{A}, \mathbf{S}^m\right) \tag{34}$$

$$\mathscr{L}_S = \sum_u \Phi\left(\mathbf{A}, \mathbf{S}^u\right) \tag{35}$$

where $\Phi$ is the MSE loss, and $\mathbf{A}$ is the adjacency matrix of SL graph.

In addition, because different data sources may play various roles in SL prediction, the author further designs an edge-level attention mechanism to assign different weights to support views to merge all rebuilt graphs.

Based on the attention scheme, a weighted similarity matrix $\mathbf{W}_{supp}$ can be derived as

$$\mathbf{W}_{supp} = \sum_u^{n_s} a^u \odot \mathbf{S}^u \tag{36}$$

where $\odot$ is the element-wise multiplication and $a^u$ is the normalized weight for the support view $u$.

And then, the final score matrix $\mathbf{S}_P$ will be derived by combining the main view matrix $\mathbf{S}_m$ and the weighted score matrix $\mathbf{W}_{supp}$ from the support views as follows:

$$\mathbf{S}^P = \mathbf{S}^m + c\mathbf{W}_{supp} \tag{37}$$

where $c$ is a hyper-parameter to control the contribution of $\mathbf{W}_{supp}$ for final prediction.
Therefore, the final prediction loss $\mathscr{L}_P$ is

$$\mathscr{L}_P = \sum_u \Phi\left(\mathbf{A}, \mathbf{S}^P\right) \tag{38}$$

Combine the reconstruction loss and prediction loss, the overall loss $\mathscr{L}_{Total}$ can be obtained as follows:

$$\mathscr{L}_{\text{Total}} = \mathscr{L}_M + \alpha\mathscr{L}_S + \beta\mathscr{L}_P \tag{39}$$

where $\alpha$ and $\beta$ are hyper-parameters, controlling the contributions from $\mathscr{L}_S$ and $\mathscr{L}_P$, respectively.

##### 2.5 PTGNN

Long et al. introduced a new Pre-Training Graph Neural Networks-based framework named PT-GNN[19], which is utilized to integrate different data sources of link prediction in biomedical networks. Protein sequences contain rich knowledge, and CNNs are able to learn high-order protein features from their sequences for various applications. Following Lee et al.[20] and Nguyen et al.[21], the authors adopt CNN to encode protein features from sequence data. In particular, they split a protein sequence $s_p$ into a set of overlapping $n$-gram amino acid segments with $r$ as the size of sliding window. Assume that a $n$-gram amino acid segment is considered as a word, and each word is represented by a $d_1$-dimension feature vector. The feature vectors of all words are denoted by $\mathbf{F}_w \in \mathbb{R}^{N_w \times d_1}$, where $N_w$ denotes the number of all possible words in the dataset, and each row of $\mathbf{F}_w$ is the feature vector of a possible word. The word feature matrix $\mathbf{F}_w$ is set as a trainable parameter matrix, which is randomly initialized and can be updated in the pre-training phase for more accurately capturing the intrinsic features of sequences.

Using a CNN-based sequence encoder, an initial feature matrix $\mathbf{X}_p \in \mathbb{R}^{N_w \times d_1}$ can be obtained. Subsequently, the authors propose a GCN-based interaction graph encoder to learn protein representations with input graph $\mathscr{G}_p$ and initial node feature $\mathbf{X}_p$. For a node $v_i$ in $\mathscr{G}_p$, the main purpose of the graph encoder is to learn its representation by iteratively aggregating the representations of its neighbors. The $\ell$-th layer of a GCN-based graph encoder is as follows:

$$\mathbf{H}_p^{(\ell)} = \text{ReLU}\left(\tilde{\mathbf{A}}_p \mathbf{H}_p^{(\ell-1)} \mathbf{W}_2^{(\ell-1)} + \mathbf{b}_2^{(\ell-1)}\right) \tag{40}$$

where $\tilde{\mathbf{A}}_p$ is the normalized diagonal adjacency matrix with self-connection, $\mathbf{H}_p^{(\ell-1)}$ denotes the outputs of the model at the $(\ell-1)$-th layer, which $\mathbf{H}_p^{(0)}$ is defined as the input feature matrix $\mathbf{X}_p$. $\mathbf{W}_2^{(\ell-1)}$ and $\mathbf{b}_2^{(\ell-1)}$ are trainable weight matrix and bias vector, respectively. After several GCN layers, the representations of proteins $\mathbf{H}_p \in \mathbb{R}^{N_p \times d_3}$ can be adopted, where $d_3$ denotes the dimension of the protein representations. Similarly, the representations of proteins $\mathbf{H}_g \in \mathbb{R}^{N_p \times d_3}$ from the protein GO graph $\mathscr{G}_g$ can be obtained with node features $\mathbf{X}_p$.

The prediction scores and loss functions of SL are as follows:

$$\mathbf{P} = \text{ReLU}\left(\mathbf{H}\mathbf{H}^\top\right) \tag{41}$$

$$\mathscr{L} = \sum_{(i,j)\in\Omega^+\cup\Omega^-} \Phi(\mathbf{P}(i,j), \mathbf{A}(i,j)) + \delta\|\Theta\|_F^2 \tag{42}$$

where $\mathbf{P}$ is the reconstructed score matrix where each element describes the interaction score for a node pair, $\mathbf{H} = \lambda\mathbf{H}_p + (1-\lambda)\mathbf{H}_g$ is the final representations of proteins obtained from the GCN-based graph encoder, $\lambda$ a tuning parameter which controls the influence of the representations from different data, $\Theta$ is the parameter matrix of the pre-training model. $\delta$ is weight factor that is used to control the influence of $\Theta$, $\Phi(\cdot)$ is the MSE loss.

## 2.6 KG4SL

The knowledge graph (KG) integrates multi-source data through standardized semantics, and can support the mining and analysis of complex associated data through the map reasoning ability. Therefore, Wang et al. proposed a GNN based model, KG4SL[22], which integrates the KG into the task of SL prediction. The model is basically formed of three sections.

**Gene-specific weighted subgraph.** Given an SL-related gene, a weighted subgraph from KG can be constructed. Identifying relevant nodes and determining the weights are two key operations to construct the gene-specific weighted subgraph. For gene $g$, $N(g)$ denotes the set of neighbors of $g$, from which $k$ neighbors are screened out and denoted as $P(g)$. In a gene-specific subgraph, different weights for edges can be assigned to describe the importance of the relations. For an SL pair $(g_i, g_j)$, the weight for an edge $r_{g,g'}$ in $g_i$'s subgraph is computed by $\omega^{g_i}_{g,g'} = g\left(\mathbf{g}_j, \mathbf{r}_{g,g'}\right)$, where $g$ is one of the entities in the subgraph of $g_i$, and $g' \in P(g)$. $\mathbf{g}_j$ and $\mathbf{r}_{g,g'}$ are the feature embedding of gene $g_j$ and relation $r_{g,g'}$, respectively. $g$ is an inner product function.

**Aggregation of node representations.** For any central entity $e$ in the subgraph of gene $g_i$, the representations of all its picked neighbors were aggregated to update its own representation. To show the topological neighborhood structure of entity $e$ in the KG, the weighted average combination of $e$'s neighborhood is computed as follows:

$$\mathbf{e}_{P(e)} = \sum_{e' \in P(e)} \tilde{\omega}^{e_i}_{e,e\prime}\mathbf{e}' \tag{43}$$

where $\mathbf{e}$ is the representation of entity $e$, gene $g_i$ and gene $g_j$ are a pair in the SL matrix, and $\tilde{\omega}^{e_i}_{e,e} = \frac{\exp\left(\omega^{e_i}_{e,e}\right)}{\sum_{\hat{e} \in P(e)} \exp\left(\omega^{e_i}_{e,\hat{e}}\right)}$ is the normalized gene-relation score by applying a softmax function.

After obtaining the picked neighbors' representation $\mathbf{e}_{P(e)}$ of a central entity in one hop, similar to Wang et al.[23], it integrates the entity representation $\mathbf{e}$ in to a single vector to update $\mathbf{e}$:

$$\mathbf{e}[h+1] = \phi\left(\mathbf{W}\left(\mathbf{e}[!h] + \mathbf{e}_{P(e)}\right) + \mathbf{b}\right) \tag{44}$$

where $\mathbf{W}$ and $\mathbf{b}$ are the linear transformation weight and bias, respectively, and $\phi$ is an activation function. After aggregating neighbors' information through $H$ hops, the final feature representation of gene $\hat{\mathbf{g}}_i$ is $\mathbf{e}[!h]$, $\hat{\mathbf{g}}_j$ is obtained in the same way.

**SL prediction.** Finally, the predicted interaction probability between gene $g_i$ and gene $g_j$ is calculated by $\hat{s}_{i,j} = \phi\left(f\left(\hat{g}_i, \hat{g}_j\right)\right)$, where $f$ is the inner product function and $\phi$ is a sigmoid function, squashing the output to a range between 0 and 1.

The base loss $L$ is computed through cross-entropy of the truth label and the predicted label for the edges, represented as follows:

$$L = \max\left(\hat{s}_{i,j}, 0\right) - \hat{s}_{i,j} \times s_{i,j} + \log\left(1 + \exp\left(-\left|\hat{s}_{i,j}\right|\right)\right) \tag{45}$$

where $\hat{s}_{i,j}$ is the predicted label and $s_{i,j}$ is the truth label for the edge.

With an L2-regulatizer, the final loss is defined as:

$$\min_{\mathbf{W},\mathbf{A},\mathbf{b}} \ell = \min_{\mathbf{W},\mathbf{A},\mathbf{b}} \sum_{i,j} J + \alpha\|\Gamma\| \tag{46}$$

where $\|\Gamma\| = \frac{\|\mathbf{e}\| + \|\mathbf{r}\| + \|\mathbf{W}\|}{2}$, $\mathbf{A}$ is the trainable weight matrix in which each element represents the gene-relation score and $\alpha$ is a balancing hyper-parameter.

#### 2.7 PiLSL

Liu et al. proposed PiLSL[24], a GNN model based on paired interactive learning, to learn the paired interactive representation of two genes with SL interaction. The author claims that most of the current methods are to learn the expression of each gene independently, ignoring the representation of the pairwise interaction between two genes. PiLSL consists of three steps to learn the representations of pairwise interactions.

**Pairwise enclosing graph extraction.** There are abundant biomedical entities connected to the genes by various relations in SynLethKG and there exists interaction information between two genes, authors focus on the local enclosing graph of each gene pair to capture the gene-gene interaction information. For a given gene pair $u$ and $v$, the enclosing graph is defined as $\mathscr{G}_{\mathbf{en}} = \{(u,r,v) \mid u,v \in \mathscr{N}_k(u) \cap \mathscr{N}_k(v), r \in \mathscr{R}\}$, where $\mathscr{N}_k(u) = \{s \mid d(s,u) \le k\}$ and $\mathscr{N}_k(v) = \{s \mid d(s,v) \le k\}$, $d(\cdot,\cdot)$ denotes the shortest distance between two elements.

**Attentive embedding propagation.** To reduce the noisy information and extract biological meaning from the prediction, attentive embedding propagation was adapted to discriminate the importance among the edges in the enclosing graph, which consists of three components: message passing, relation-aware attention and message aggregation.

At the $l$-th layer, message passing used to combine the representations of the neighbors with the node $u$,

$$\mathbf{m}_{\mathscr{N}_u}^{(l)} = \sum_{r=1}^{N_R} \sum_{v \in \mathscr{N}_r(u)} a_{r(u,v)}^{(l)} \mathbf{W}_r^l \mathbf{h}_v^{(l-1)} \tag{47}$$

where $N_R$ is the total number of the relations in SynLethKG, $\mathscr{N}_r(u)$ denotes the set of directly connected neighbors of node $u$ under relation $r$, $\mathbf{W}_r^l$ is the weighted matrix to transform hidden representation in the $l$-th layer over relation $r$, $a_{r(u,v)}^{(l)}$ is the attention weight, controlling how much message being passed from $v$ to $u$ via relation $r$.

Then, the node representations are updated by node self-representation and message passing from its neighbors as follows:

$$\mathbf{h}_u^{(l)} = \mathrm{ReLU}\left(\mathbf{W}_{\text{self}}^{(l)} \mathbf{h}_u^{(l-1)} + \mathbf{m}_{\mathscr{N}_u}^{(l)}\right), \tag{48}$$

where $\mathbf{W}_{\text{self}}^{(l)}$ is the weight matrix of transforming the node self-representation, $\mathbf{h}^{(l)}$ is the embeddings of all entities in the layer $l$, $\mathbf{h}^{(0)}$ were initialized by TransE.

**Latent and explicit features fusion.** Latent features consist of information at the node level and the graph level. At the node level, the representation $\mathbf{h}^{(l)}$ will be obtained by attentive embedding propagation. At the graph level, the representation $\mathbf{h}_{\mathscr{G}_{\mathbf{en}}}^{(l)}$ can be got by the average of all node representations in the enclosing graph. The representation of $\mathbf{h}_{\mathscr{G}_{\mathbf{en}}}^{(l)}$ at layer $l$ is:

$$\mathbf{h}_{\mathscr{G}_{\mathbf{en}}}^{(l)} = \frac{1}{|\mathscr{V}|} \sum_{i \in \mathscr{V}} \mathbf{h}_i^{(l)} \tag{49}$$

where $\mathscr{V}$ is the set of nodes in the enclosing graph $\mathscr{G}_{\mathbf{en}}$.

The final predicted interaction probability is $\hat{p}_{uv} = \mathbf{W}_{pred}\mathbf{h}_{uv}$, where $\mathbf{W}_{pred}$ is the weight matrix of the decoder.

Two kinds of losses are used for optimization:

$$L_{CE} = -\frac{1}{|\tau|} \sum_{(u,v)\in\tau} p_w \log \hat{p}_{uv} + (1 - p_{uv}) \log(1 - \hat{p}_{uv}) \tag{50}$$

$$L_W = \|\mathbf{W}_{all}\|_2^2 \tag{51}$$

where $\tau$ is the sampled training set of gene pairs and $p_{uv}$ denotes true label for the SL interaction, $\mathbf{W}_{all}$ denotes all weight parameters.

The total loss is $L_{Total} = L_{CE} + \frac{\lambda}{2} L_W$, where $\lambda$ is the coefficient of $L_W$.

##### 2.8 NSF4SL

Because the prediction task of SL usually encounters the problem of lacking high-quality negative samples, inspired by the negative-sample-free contrastive learning frameworks, Wang et al. proposed a new SL prediction model, NSF4SL[25], which gives priority to the positive samples. NSF4SL first extracts the feature augmentation view of each gene, and then inputs the augmentation features and original features of two genes in the SL gene pair into two branches of the contrastive learning framework respectively. First, they extract a feature-wise augmented view for each gene. Second, they feed augmented features of the two genes in an SL pair into the online branch, and the original features of them into the target branch. Third, they obtain the gene embeddings and maximize the similarity between them.

**Feature-wise data augmentation** is designed to address the sparsity of the training data. Given a gene training set $\mathscr{G}_t$, for a gene $g \in \mathscr{G}_t$, they obtain the initial feature of $g$ as $\mathbf{x}_g = \{x_g^{(1)}, \cdots x_g^{(d)}\}$, where $x_g^i$ denotes the $i^{th}$ element of $\mathbf{x}_g$, and $d$ denotes the dimension of the vector. Then, they select a random subset of $\mathbf{x}_g$ and replace them by the respective feature-level average values. Specifically, in each iteration of training, they sample an augmented view $\mathscr{V}_g$ for each gene $g$ of length $q$,

$$\mathscr{V}_g = u(\mathbf{x}_g), \qquad q = \lfloor r \cdot d \rfloor, \tag{52}$$

where $u(\cdot)$ is the uniformly sampling function, $\lfloor \cdot \rfloor$ is the floor function and $r$ is the defined augmented ratio. Next, they obtain the augmented representation $\widetilde{\mathbf{x}}_g$ for gene $g$ as follows,

$$\widetilde{x}_g^{(i)} = \begin{cases} \bar{x}^{(i)}, & \text{if } x_g^i \in \mathscr{V}_g \\ x_g^{(i)}, & \text{otherwise} \end{cases} \tag{53}$$

where $\bar{x}^{(i)}$ is mean of the $i^{th}$ element over the whole gene set $\mathscr{G}_t$.

**Negative-sample-free architecture** is to avoid the negative sampling bias in the models for SL prediction. They employ a non-symmetric two-branch network as the backbone model. Instead of classifying SL and non-SL pairs, this network learns to recognize similarities between input SL pairs. The architecture relies on two parallel branches: they contain an online encoder $f_\theta$ and a target encoder $f_\xi$ with the corresponding trained weights $\theta$ and $\xi$, respectively. $f_\theta$ and $f_\xi$ share the same architecture but have different weights. One key design is the predictor $p_\varphi$ following the online encoder. As the target encoder is used to output the target SLs, $p_\varphi$ uses the gene embeddings of the same SLs from the online encoder to be close to them.

In the end of training, they adopt contrastive learning loss to maximize the similarity between the representations of two genes in an SL pair. Let $(\mathbf{g_i}, \mathbf{g_j})$ be the features of an input pair of genes known to have SL relation with each other. The online branch output are

$$\mathbf{g}_i^o = p_\varphi\left(f_\theta(\widetilde{\mathbf{g}}_i)\right) \quad \text{and} \quad \mathbf{g}_j^o = p_\varphi\left(f_\theta(\widetilde{\mathbf{g}}_j)\right), \tag{54}$$

and the target branch output as

$$\mathbf{g}_i^t = f_\xi(\mathbf{g}_i) \quad \text{and} \quad \mathbf{g}_j^t = f_\xi(\mathbf{g}_j). \tag{55}$$

They minimize the following objective function $\mathscr{L}$ to maximize the similarity between the representations of the two genes from the two branches,

$$\mathscr{L} = -\frac{1}{2}\left(h(\mathbf{g}_i^o, \mathbf{g}_j^t) + h(\mathbf{g}_j^o, \mathbf{g}_i^t)\right), \tag{56}$$

where $h(\cdot)$ is the inner product function between two representations.

**Table S1.** List of classic machine learning method for SL prediction.

| Model & Ref. | Year | Description |
|---|---|---|
| Paladugu et al.[26] | 2008 | A SVM system that uses graph-theoretic properties of two proteins in a protein interaction network as input features for prediction of synthetic sick/lethal interactions. |
| Pandey et al.[27] | 2010 | The author defines a large number of more comprehensive SL independent features, and uses these features to develop a MNMC framework for predicting SL interaction. The system can realize multiple classification processes at the same time. |
| MetaSL[28] | 2014 | The author proposed an ensemble method for predicting yeast SL, named MetaSL, which integrates the output of 17 genomic and proteomic features and 10 classification methods. |
| Yin et al.[29] | 2019 | The author developed a DT based method and used the features based on mutation and copy number variation level to predict gene pairs with SL interaction. |
| DiscoverSL[30] | 2019 | This is a multi-parameter RF classifier, to predict and visualize synthetic lethality in cancers using multi-omic cancer data. |
| Li et al.[31] | 2019 | Using the selected functional features obtained from GO and KEGG data, the author proposed a method based on RF to predict SL gene pairs. |
| SLant[32] | 2019 | This is an RF based method, which works by identifying and exploiting conserved patterns in protein interaction network topology both within and across species. |
| Wu et al.[33] | 2021 | Multiple types of gene similarity measures are integrated and k-NN algorithm is applied to achieve the similarity-based classification task between gene pairs. |
| De Kegel et al.[34] | 2021 | By analyzing the genome-wide CRISPR screening and the molecular map of cancer cell lines, the author identified the shared PPI and evolutionary conservation features to predict SL, and developed a classifier based on RF according to these features. |
| PARIS[35] | 2021 | PARIS is an RF based method, it predicts SL interactions by combining CRISPR viability screens with genomics and transcriptomics data across hundreds of cancer cell lines profiled within the Cancer Dependency Map. |
| SBSL[36] | 2022 | The authors introduced regularized logistic regression or RF into the selection bias resistant synthetic lethality (SBSL) prediction for each gene pair molecularly characterized from cell lines, cancer patient tissues and healthy donor tissue samples. |
| GRSMF[2] | 2019 | GRSMF is a method based on graph regularized self-representation matrix factorization (MF). It learns self-representation from known SL interactions and further integrates GO information to predict potential SL interactions. |
| CMFW[5] | 2020 | CMFW is a collective matrix factorization-based method that integrates multiple heterogeneous data sources for SL prediction. |
| $SL^2MF$[1] | 2020 | $SL^2MF$ uses logistic matrix factorization to learn gene representations, which are then used to identify potential SL interactions. The authors design an importance weighting scheme to distinguish known and unknown SL pairs and combine PPI and GO information for the prediction. |

**Table S2.** List of deep learning method for SL prediction.

| Model & Ref. | Year | Description |
|---|---|---|
| EXP2SL[37] | 2020 | A method based on semi-supervised neural network uses the cell-specific gene expression profile data obtained in the L1000 project to provide features for predicting human SL interaction. |
| DDGCN[11] | 2020 | DDGCN is the first graph neural network (GNN)-based method for SL prediction. It uses graph convolutional network (GCN) and known SL interaction matrix as features. The authors use coarse-grained node dropout and fine-grained edge dropout to address the issue of overfitting of GCNs on sparse graphs. |
| GCATSL[15] | 2021 | GCATSL proposes a graph contextualized attention network to learn gene representations for SL prediction. The authors use data of GO and PPI to generate a set of feature graphs as model inputs and introduce attention mechanisms at the node and feature levels to capture the influence of neighbors and learn gene expression from different feature graphs. |
| SLMGAE[18] | 2021 | SLMGAE is a method for predicting SL interactions by leveraging a multi-view graph autoencoder. The authors incorporate data from PPI and GO as supporting views, while utilizing the SL graph as the main view, and apply a graph autoencoder (GAE) to reconstruct these views. |
| MGE4SL[14] | 2021 | MGE4SL is a method based on Multi-Graph Ensemble (MGE) to integrate biological knowledge from PPI, GO, and Pathway. It combines the embeddings of features with different neural networks. |
| KG4SL[22] | 2021 | KG4SL is a novel model based on graphical neural networks (GNN), and the first method that utilizes knowledge graph (KG) for SL prediction. The integration of KG helps the model obtain more information. |
| PiLSL[24] | 2021 | PiLSL is a graph neural network (GNN)-based method that predicts SL by learning the representation of pairwise interaction between two genes. |
| PTGNN[19] | 2021 | PTGNN is a pre-training method based on graph neural networks that can integrate various data sources and leverage the features obtained from graph-based reconstruction tasks to initialize models for downstream link prediction tasks. |
| NSF4SL[25] | 2022 | NSF4SL is a contrastive learning-based model for SL prediction that eliminates the need for negative samples. It frames the SL prediction task as a gene ranking problem and utilizes two interacting neural network branches to learn representations of SL-related genes, thereby capturing the characteristics of positive SL samples. |
| MAGCN[38] | 2022 | A multiple attention graph convolutional network for predicting synthetic lethality. The authors used GCN to accumulate knowledge from topological structures and genetic features obtained from PPI, GO, and constructed a multi-graph attention model to learn the contributing factors of different graphs and aggregate these graphs to generate embeddings. |
| MVGCN-iSL[39] | 2023 | A multi-view graph convolutional network (GCN) model to predict cancer cell-specific SL gene pairs by integrating five biogram features and multi-omics data. |
| SLGNN[40] | 2023 | A GNN model that enables a better understanding of the underlying biological mechanisms predicting SL, which models the preferences of genes with different relationships in the knowledge graph, ultimately providing better interpretability. |

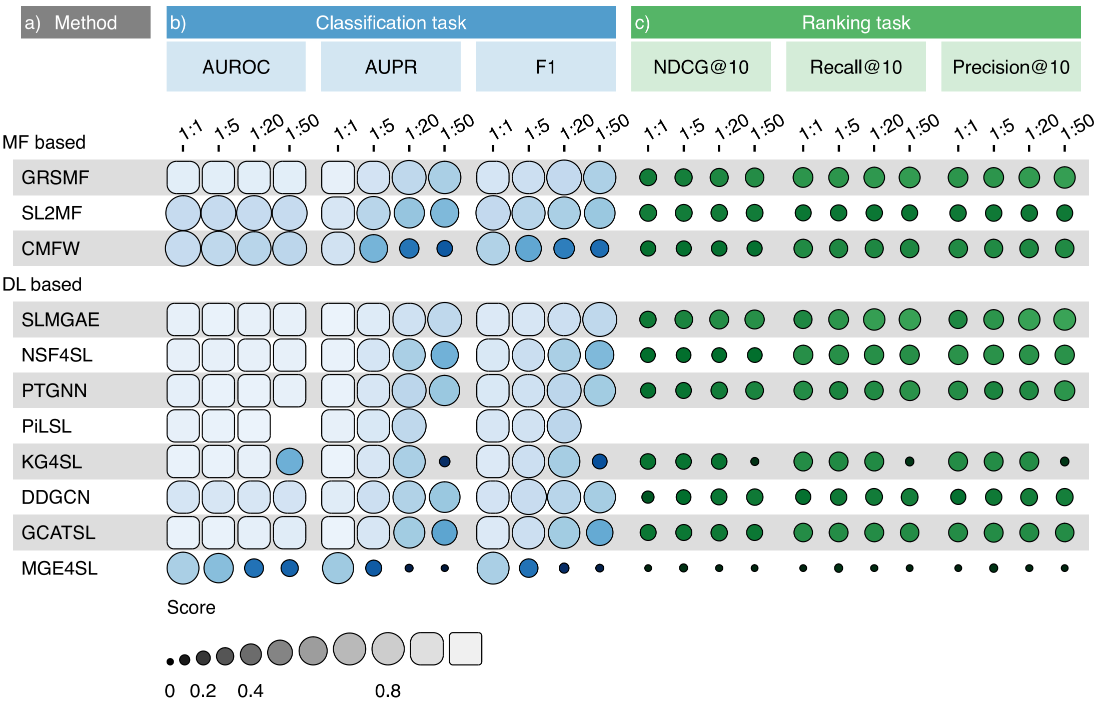


**Figure S1.** **Result under CV1 and $NSM_{Rand}$**

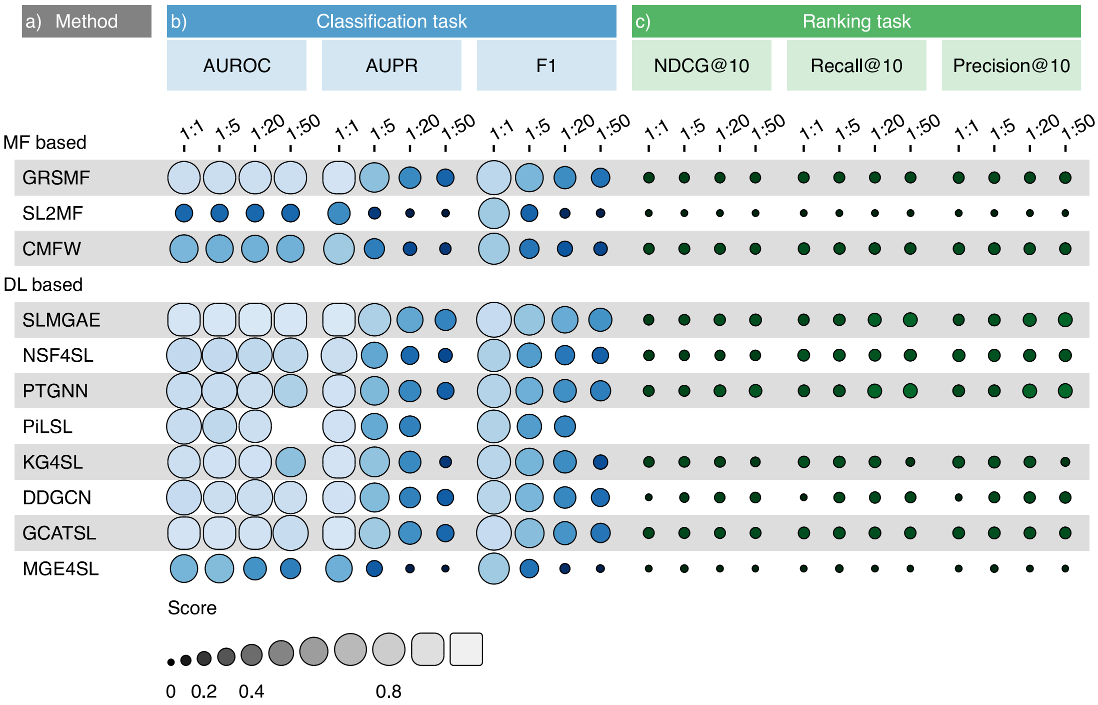


**Figure S2.** **Result under CV2 and $NSM_{Rand}$**

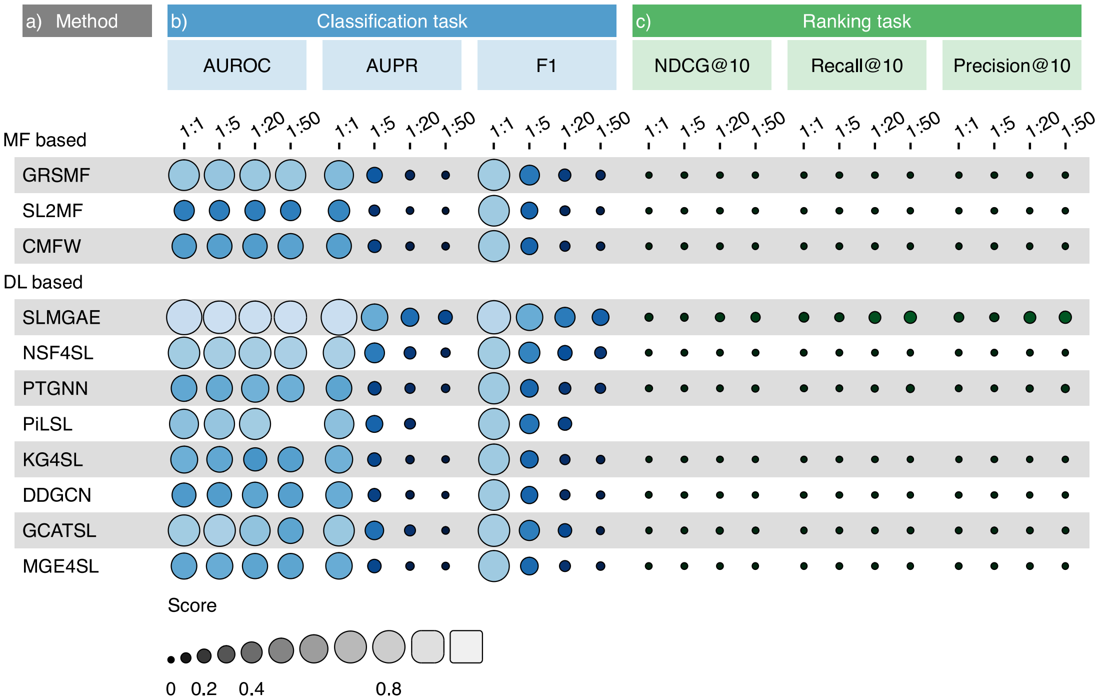


**Figure S3.** Result under CV3 and $NSM_{Rand}$

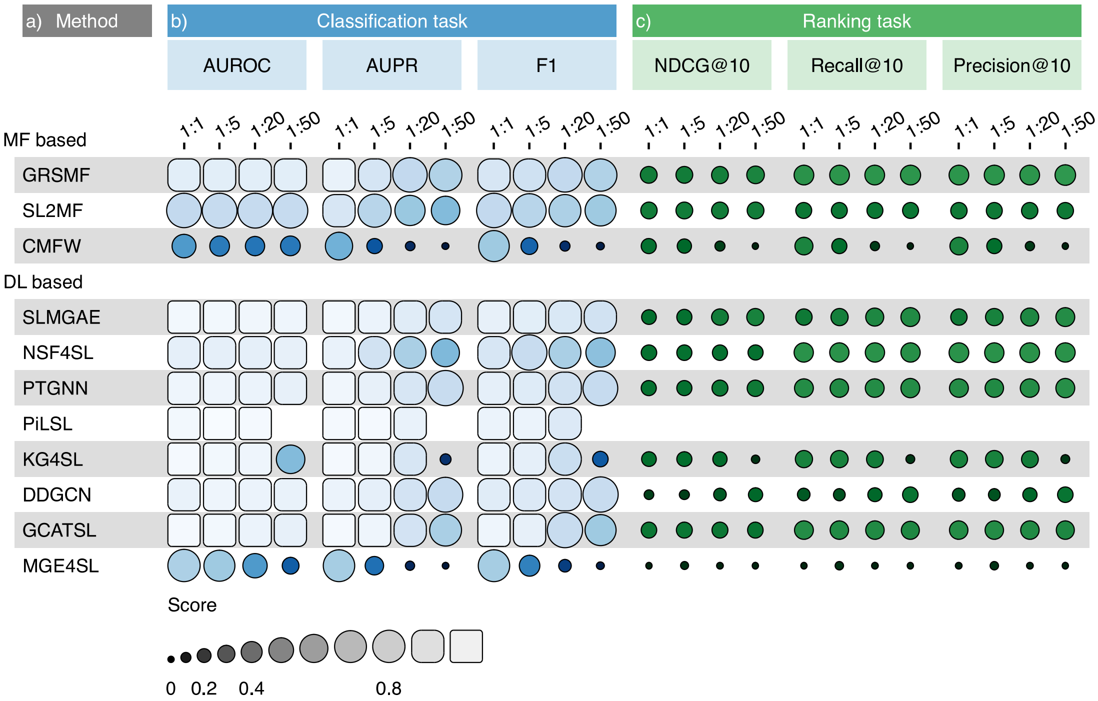


**Figure S4.** Result under CV1 and $NSM_{Exp}$

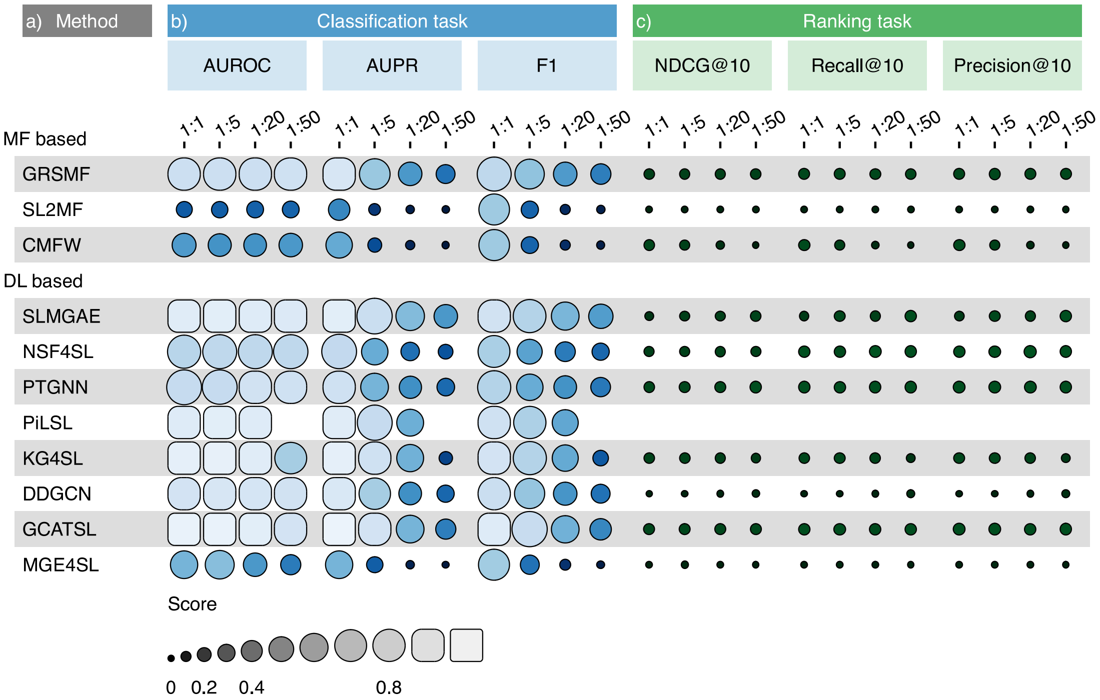


**Figure S5.** Result under CV2 and $NSM_{Exp}$

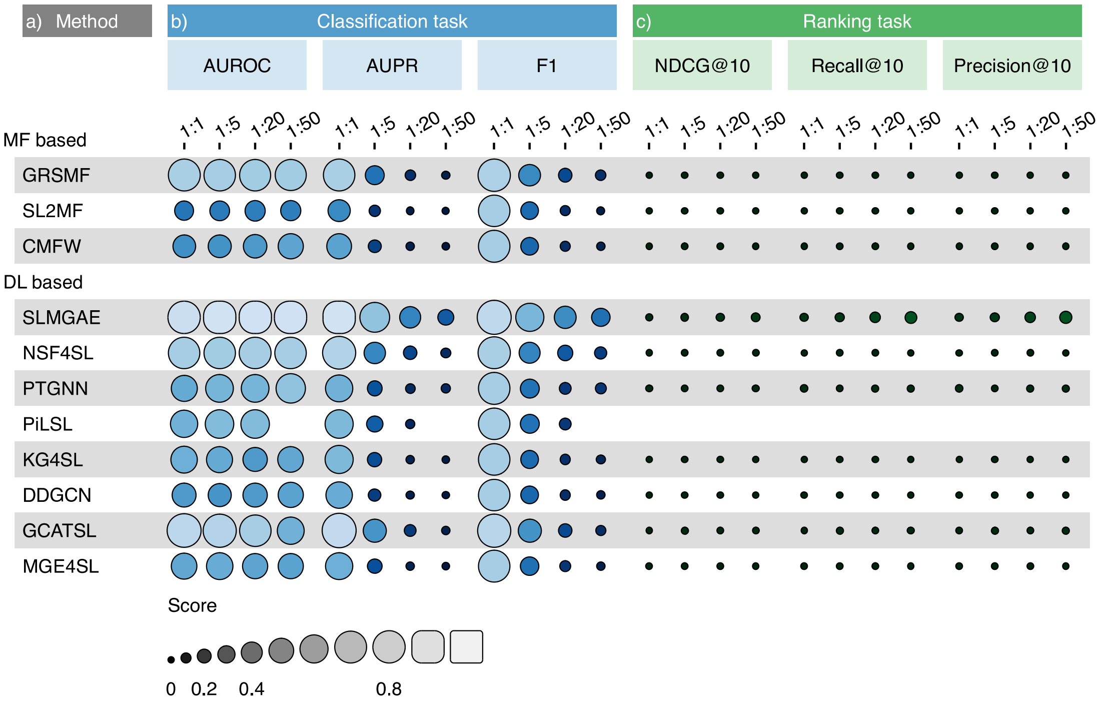


**Figure S6.** Result under CV3 and $NSM_{Exp}$

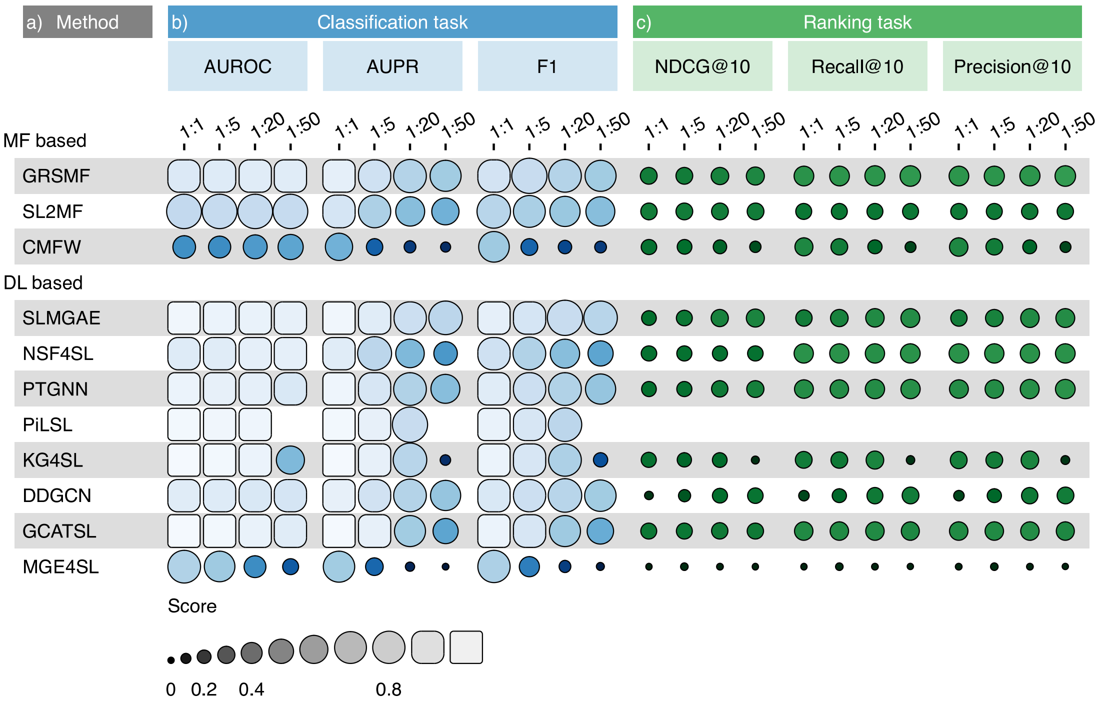


**Figure S7.** Result under CV1 and $NSM_{Dep}$

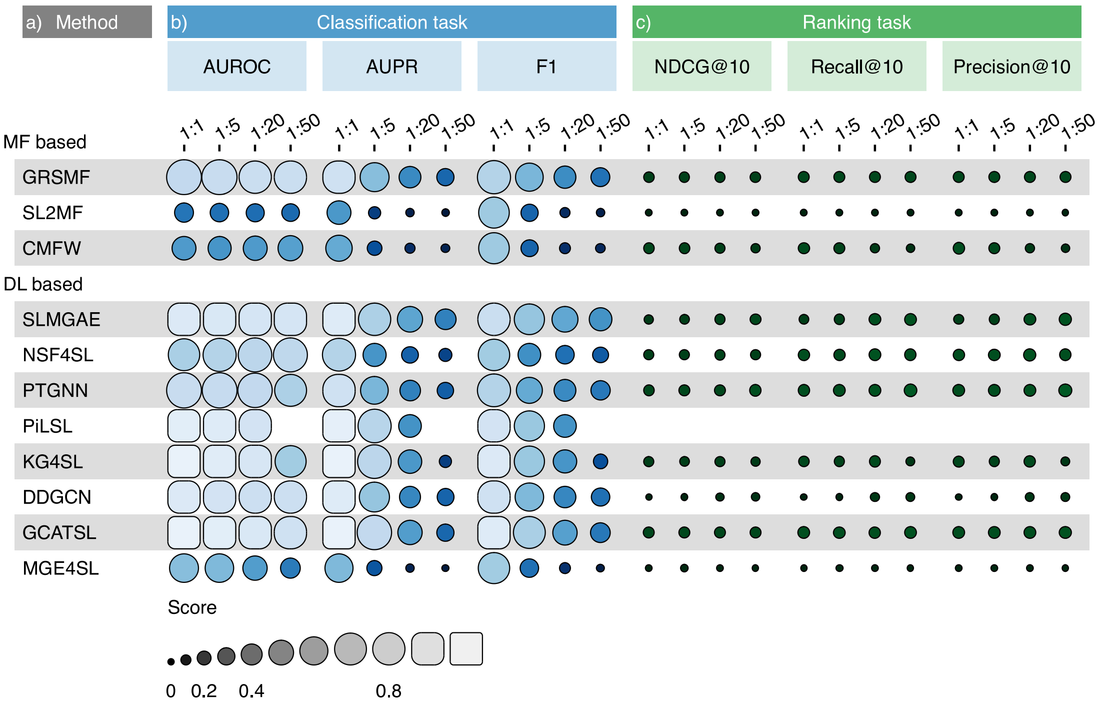


**Figure S8.** Result under CV2 and $NSM_{Dep}$

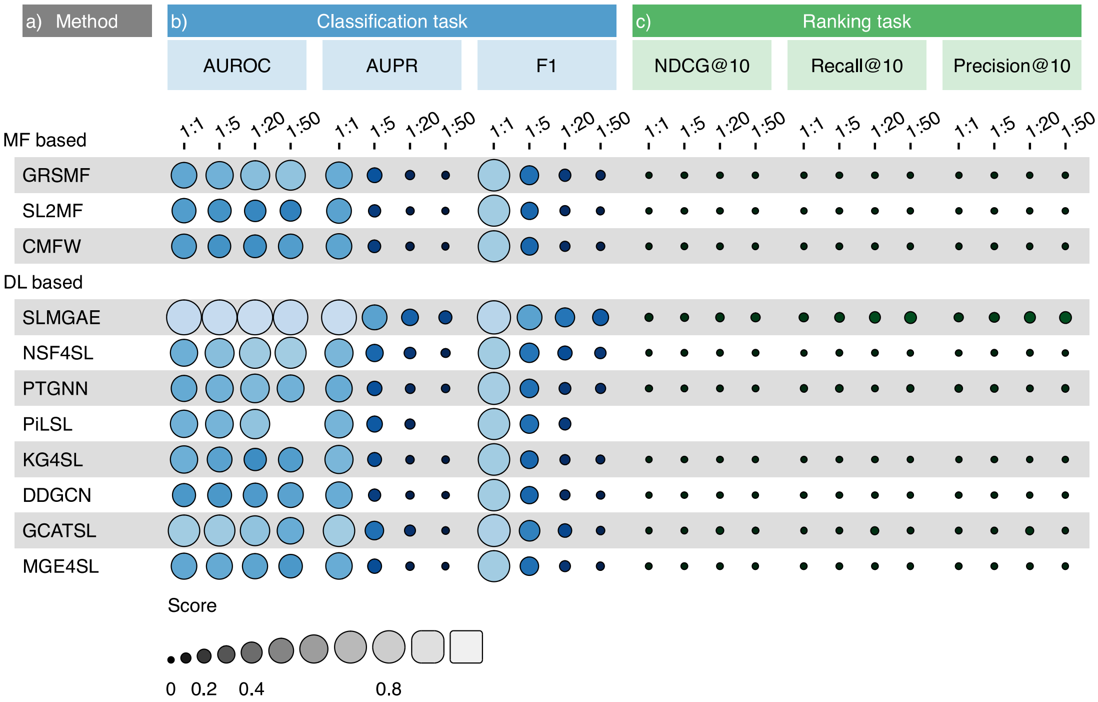


**Figure S9.** **Result under CV3 and $NSM_{Dep}$**

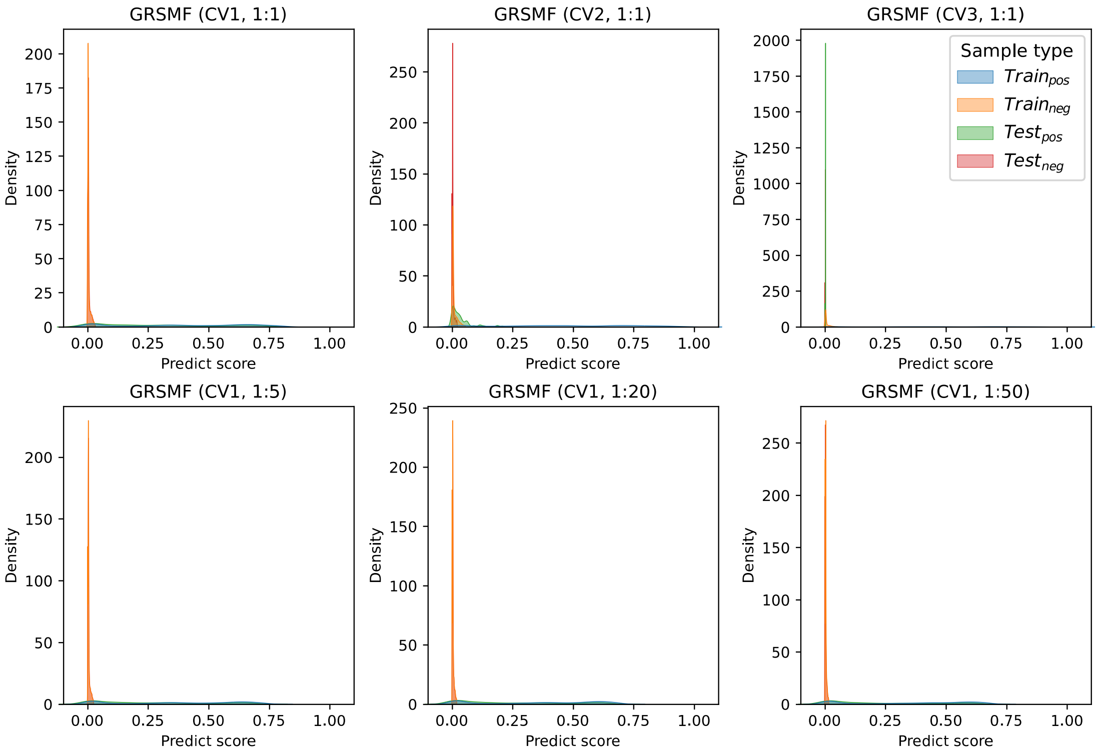


**Figure S10.** **Distribution of predicted score.** **A**, **B**, **C**, **D**, **E** and **F** show the predicted score distributions of the GRSMF model in the training and testing sets under (CV1, 1:1), (CV2, 1:1), (CV3, 1:1), (CV1, 1:5), (CV1, 1:20), and (CV1, 1:50) scenarios, respectively.

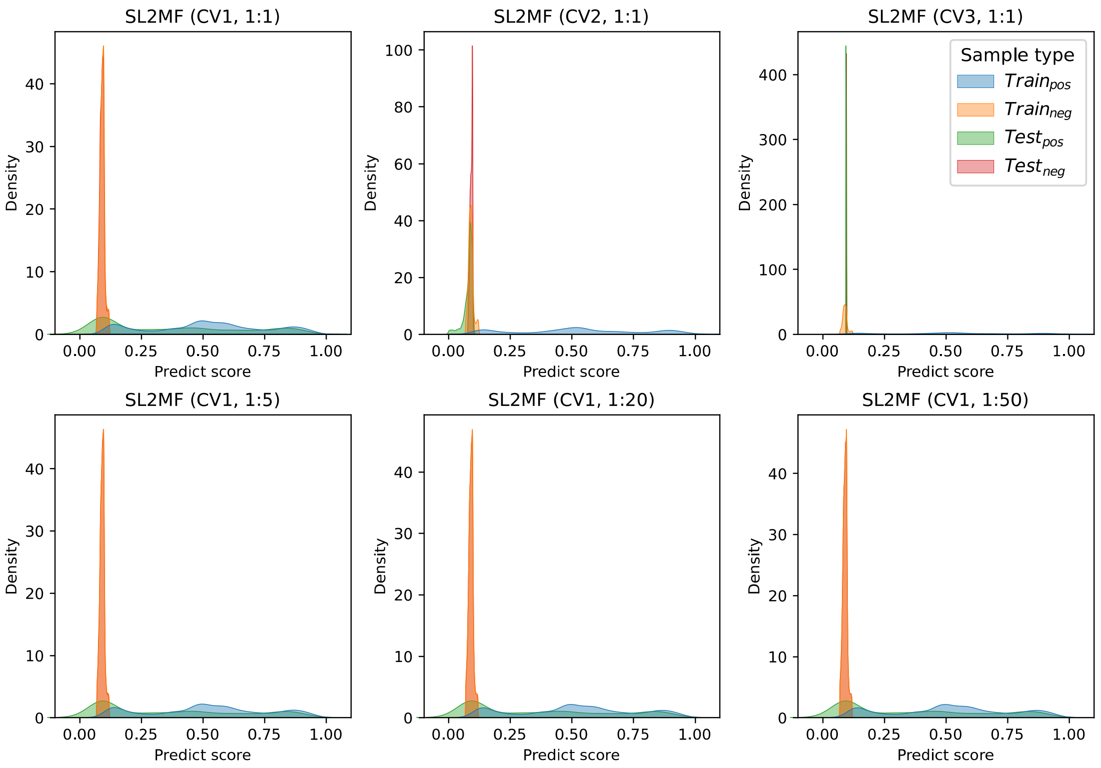


**Figure S11. Distribution of predicted score. A**, **B**, **C**, **D**, **E** and **F** show the predicted score distributions of the SL$^2$MF model in the training and testing sets under (CV1, 1:1), (CV2, 1:1), (CV3, 1:1), (CV1, 1:5), (CV1, 1:20), and (CV1, 1:50) scenarios, respectively.

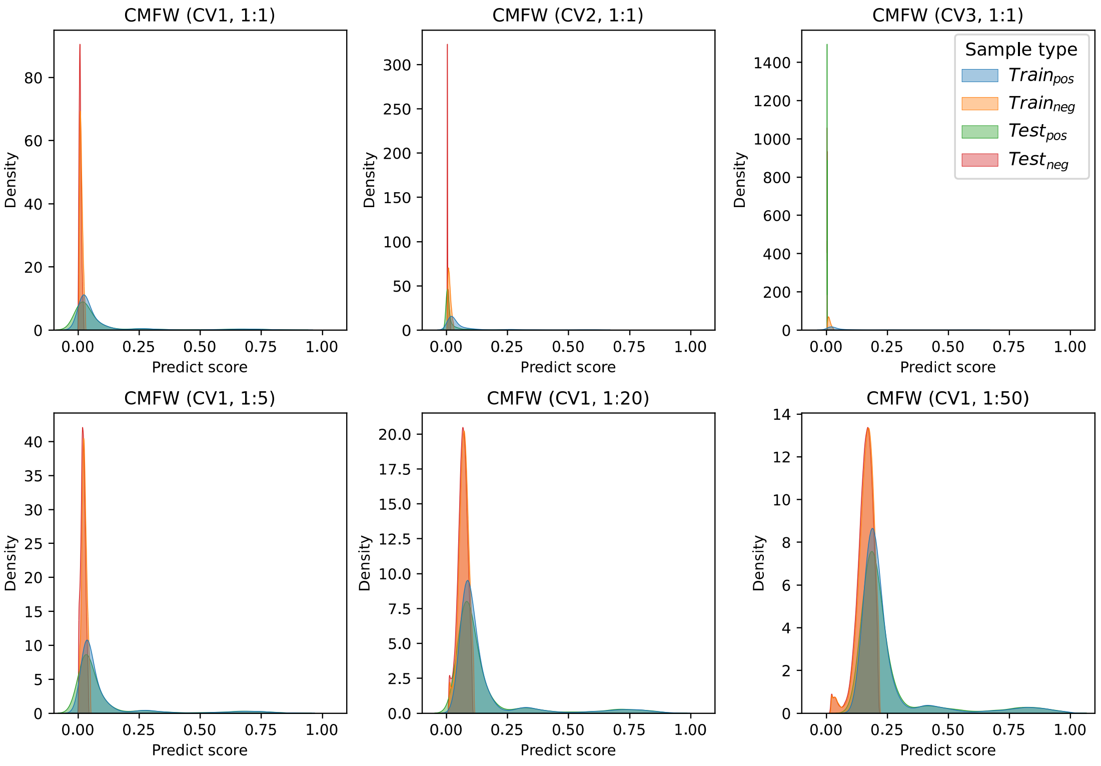


**Figure S12. Distribution of predicted score. A**, **B**, **C**, **D**, **E** and **F** show the predicted score distributions of the CMFW model in the training and testing sets under (CV1, 1:1), (CV2, 1:1), (CV3, 1:1), (CV1, 1:5), (CV1, 1:20), and (CV1, 1:50) scenarios, respectively.

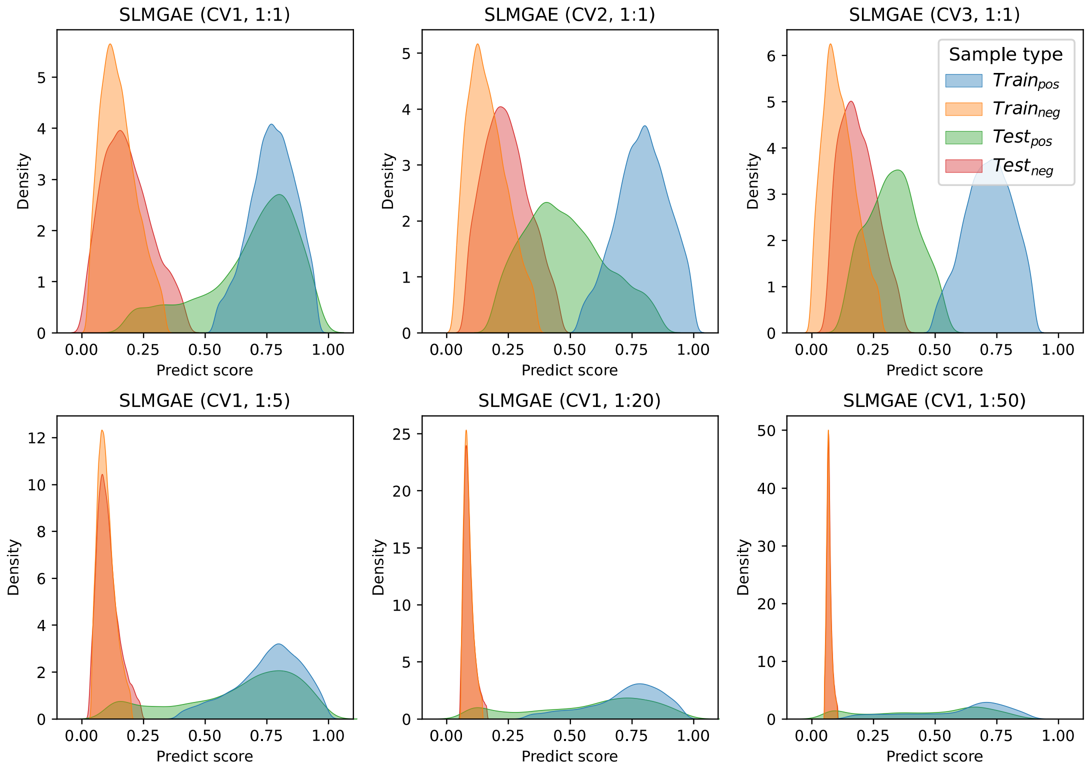


**Figure S13. Distribution of predicted score.** **A**, **B**, **C**, **D**, **E** and **F** show the predicted score distributions of the SLMGAE model in the training and testing sets under (CV1, 1:1), (CV2, 1:1), (CV3, 1:1), (CV1, 1:5), (CV1, 1:20), and (CV1, 1:50) scenarios, respectively.

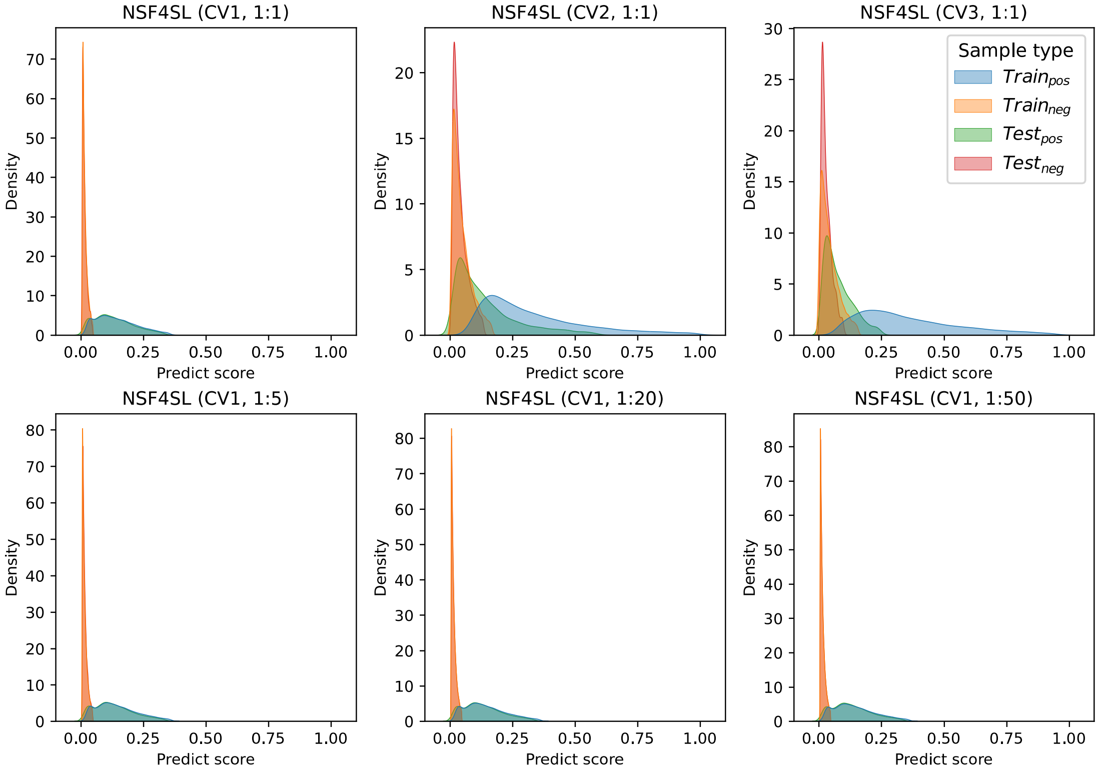


**Figure S14. Distribution of predicted score.** **A**, **B**, **C**, **D**, **E** and **F** show the predicted score distributions of the NSF4SL model in the training and testing sets under (CV1, 1:1), (CV2, 1:1), (CV3, 1:1), (CV1, 1:5), (CV1, 1:20), and (CV1, 1:50) scenarios, respectively.

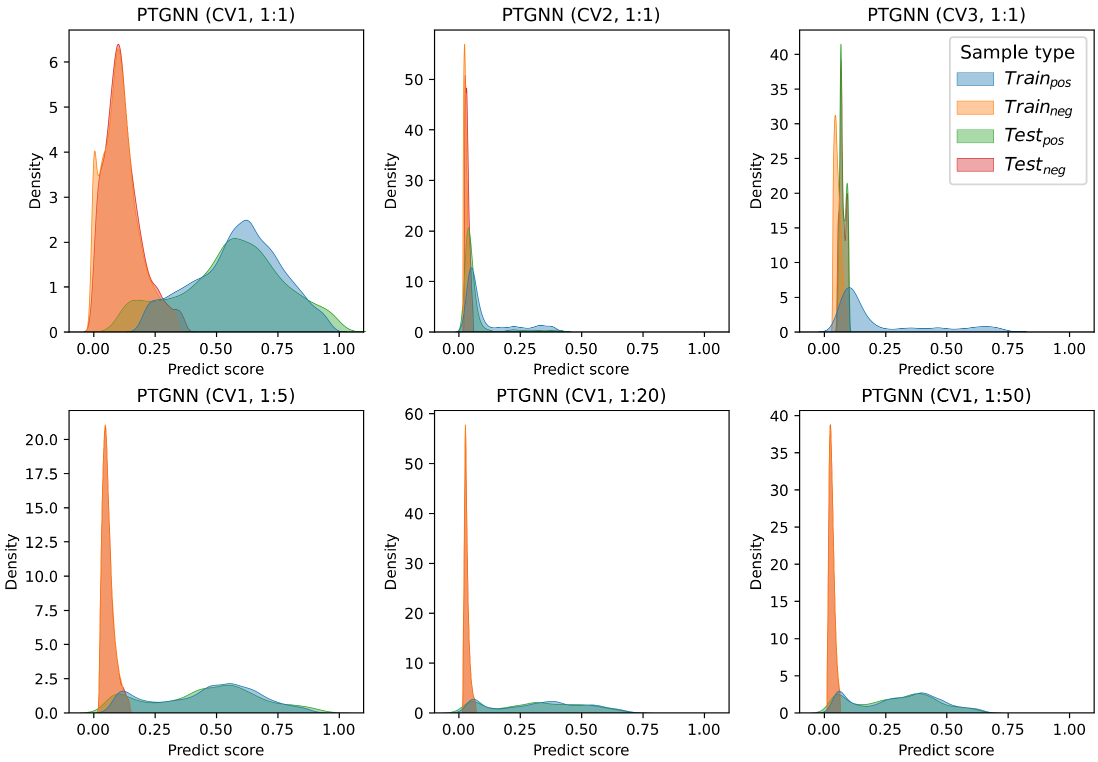


**Figure S15. Distribution of predicted score.** **A**, **B**, **C**, **D**, **E** and **F** show the predicted score distributions of the PTGNN model in the training and testing sets under (CV1, 1:1), (CV2, 1:1), (CV3, 1:1), (CV1, 1:5), (CV1, 1:20), and (CV1, 1:50) scenarios, respectively.

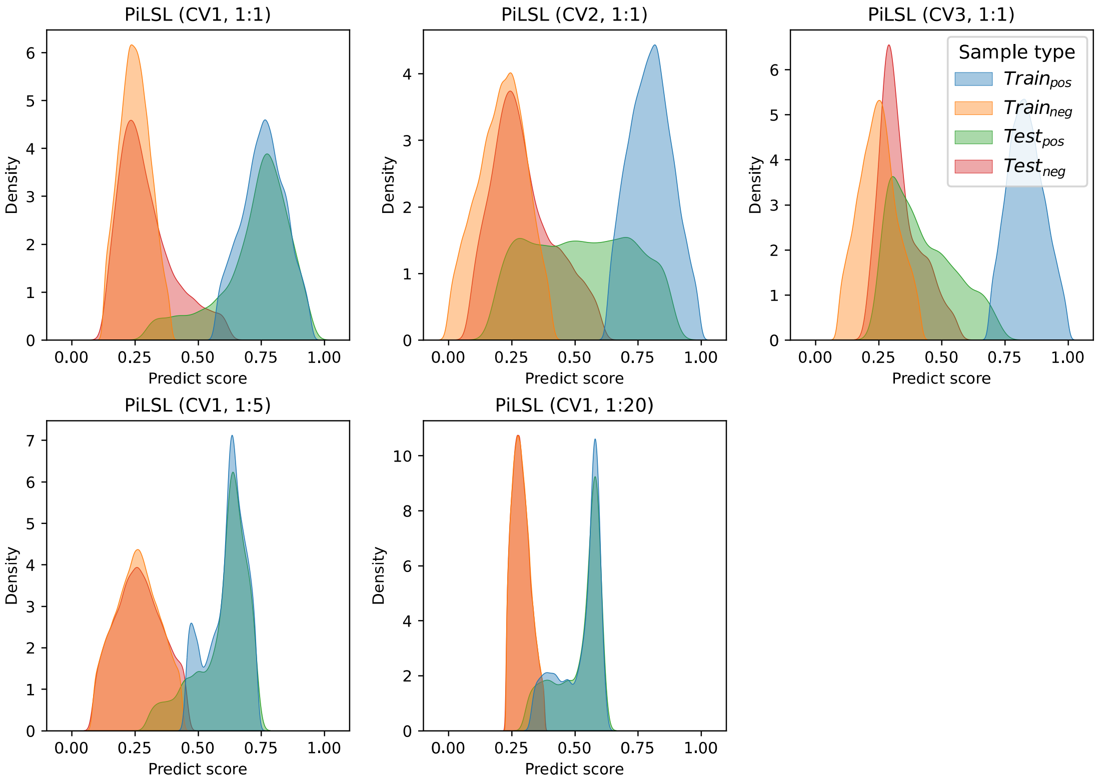


**Figure S16. Distribution of predicted score.** **A**, **B**, **C**, **D**, **E** and **F** show the predicted score distributions of the PiLSL model in the training and testing sets under (CV1, 1:1), (CV2, 1:1), (CV3, 1:1), (CV1, 1:5), (CV1, 1:20), and (CV1, 1:50) scenarios, respectively.

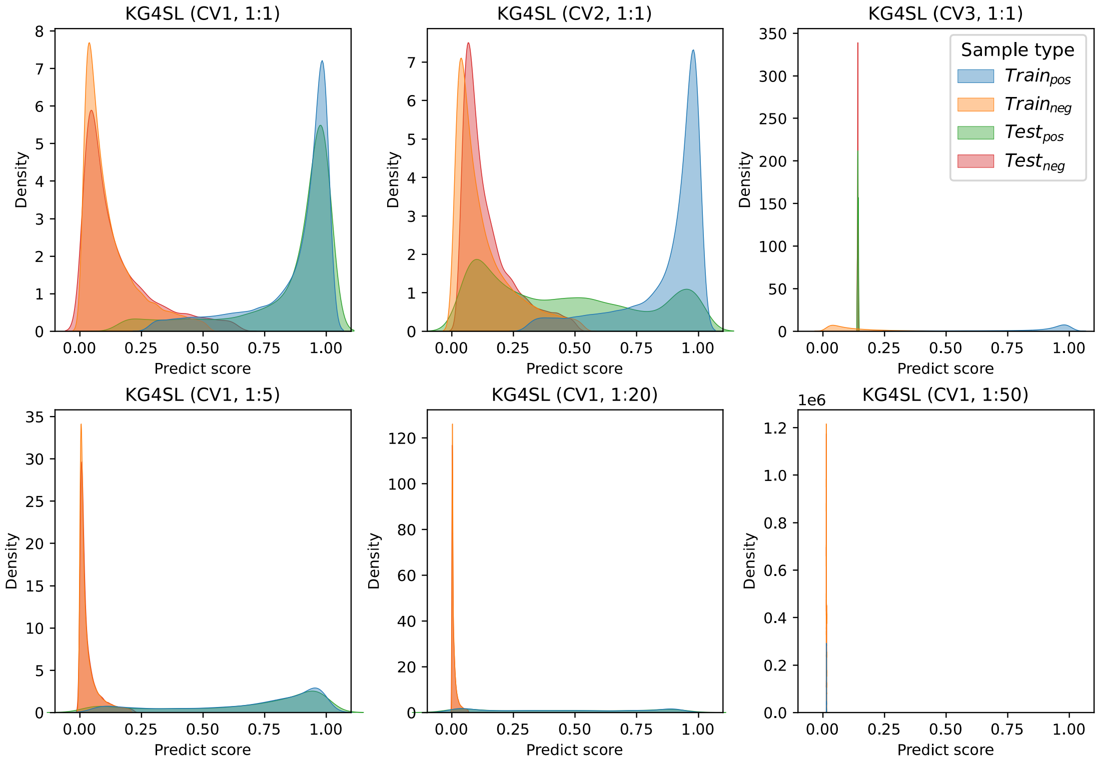


**Figure S17. Distribution of predicted score.** **A**, **B**, **C**, **D**, **E** and **F** show the predicted score distributions of the KG4SL model in the training and testing sets under (CV1, 1:1), (CV2, 1:1), (CV3, 1:1), (CV1, 1:5), (CV1, 1:20), and (CV1, 1:50) scenarios, respectively.

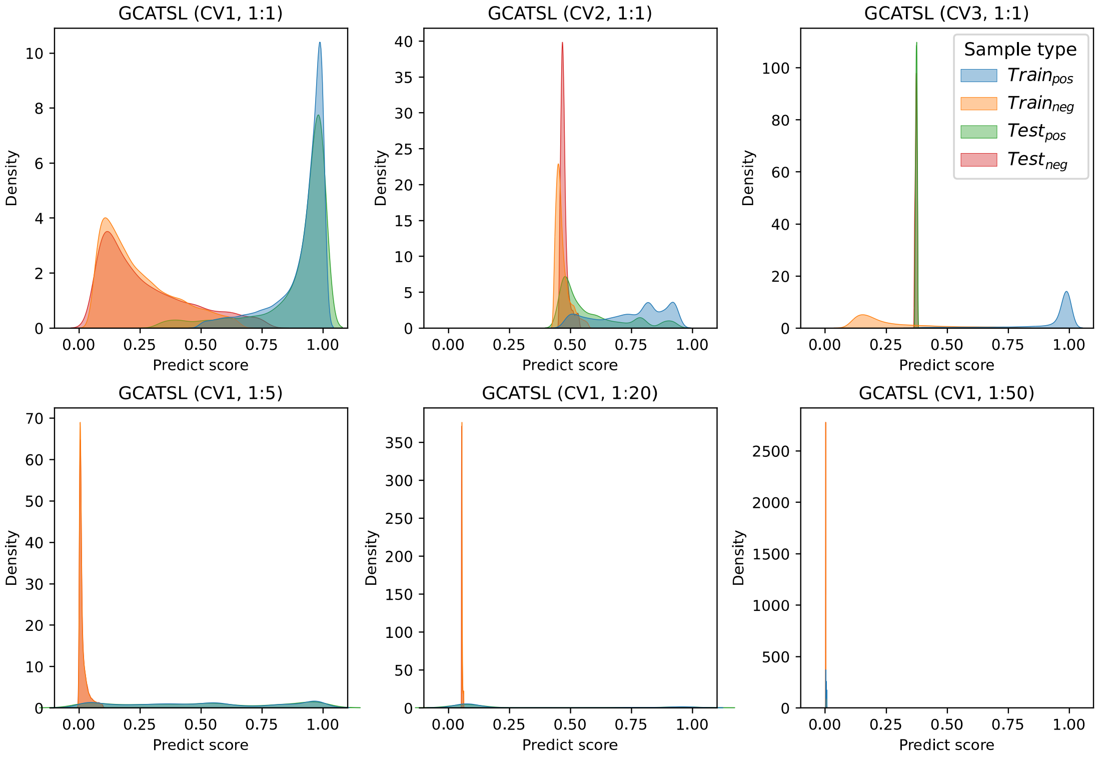


**Figure S18. Distribution of predicted score.** **A**, **B**, **C**, **D**, **E** and **F** show the predicted score distributions of the GCATSL model in the training and testing sets under (CV1, 1:1), (CV2, 1:1), (CV3, 1:1), (CV1, 1:5), (CV1, 1:20), and (CV1, 1:50) scenarios, respectively.

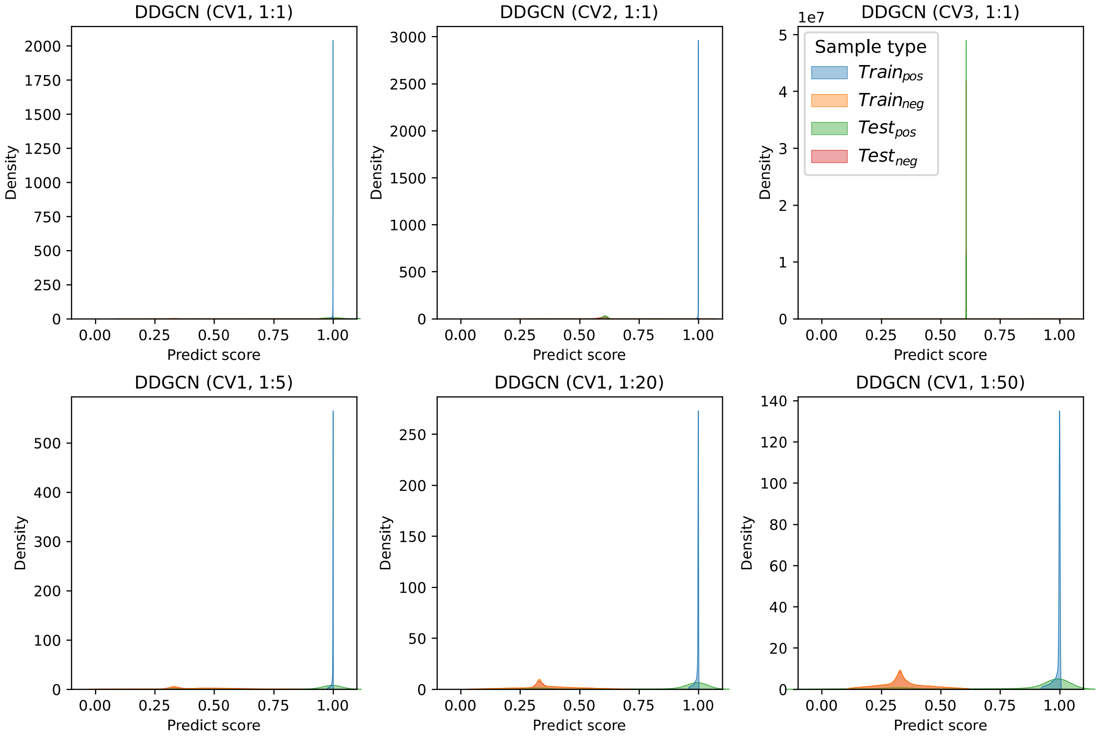


**Figure S19. Distribution of predicted score.** **A**, **B**, **C**, **D**, **E** and **F** show the predicted score distributions of the DDGCN model in the training and testing sets under (CV1, 1:1), (CV2, 1:1), (CV3, 1:1), (CV1, 1:5), (CV1, 1:20), and (CV1, 1:50) scenarios, respectively.

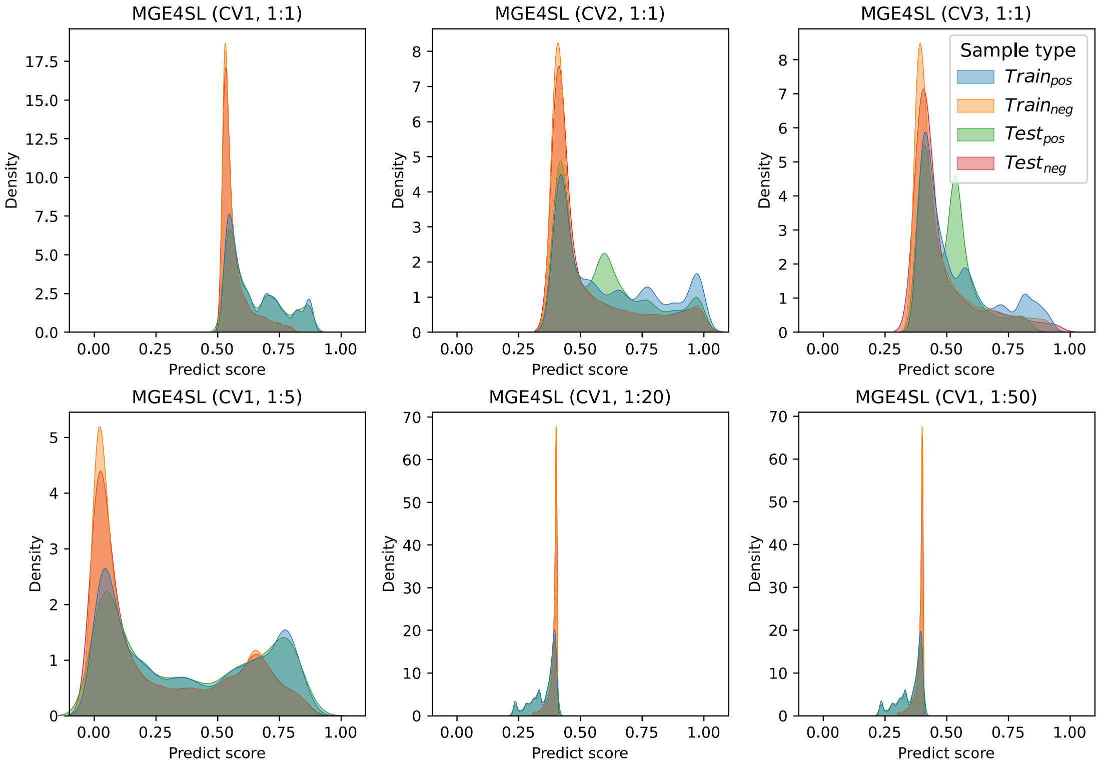


**Figure S20. Distribution of predicted score.** **A**, **B**, **C**, **D**, **E** and **F** show the predicted score distributions of the MGE4SL model in the training and testing sets under (CV1, 1:1), (CV2, 1:1), (CV3, 1:1), (CV1, 1:5), (CV1, 1:20), and (CV1, 1:50) scenarios, respectively.
